# The Ortholog Conjecture Revisited: the Value of Orthologs and Paralogs in Function Prediction

**DOI:** 10.1101/2019.12.27.889691

**Authors:** Moses Stamboulian, Rafael F. Guerrero, Matthew W. Hahn, Predrag Radivojac

**Affiliations:** Department of Computer Science, Indiana University, Bloomington, IN, U.S.A.; Department of Biology, Indiana University, Bloomington, IN, U.S.A.; Khoury College of Computer Sciences, Northeastern University, Boston, MA, U.S.A.

## Abstract

The computational prediction of gene function is a key step in making full use of newly sequenced genomes. Function is generally predicted by transferring annotations from homologous genes or proteins for which experimental evidence exists. The “ortholog conjecture” proposes that orthologous genes should be preferred when making such predictions, as they evolve functions more slowly than paralogous genes. Previous research has provided little support for the ortholog conjecture, though the incomplete nature of the data cast doubt on the conclusions. Here we use experimental annotations from over 40,000 proteins, drawn from over 80,000 publications, to revisit the ortholog conjecture in two pairs of species: (i) *Homo sapiens* and *Mus musculus* and (ii) *Saccharomyces cerevisiae* and *Schizosaccharomyces pombe*. By making a distinction between questions about the evolution of function versus questions about the prediction of function, we find strong evidence against the ortholog conjecture in the context of function prediction, though questions about the evolution of function remain difficult to address. In both pairs of species, we quantify the amount of data that must be ignored if paralogs are discarded, as well as the resulting loss in prediction accuracy. Taken as a whole, our results support the view that the types of homologs used for function transfer are largely irrelevant to the task of function prediction. Aiming to maximize the amount of data used for this task, regardless of whether it comes from orthologs or paralogs, is most likely to lead to higher prediction accuracy.

## 1 Introduction

Whole-genome sequencing of new species continues to outpace the experiments needed to annotate the function of every gene in these genomes. As a result, computational prediction of gene function is an essential tool for researchers in a range of biomedical fields. The prediction of gene function generally proceeds by the transfer of function from genes with experimental evidence to unannotated, or less-annotated, genes that are similar by some measure [42]. While several methods use multiple data types to carry out predictions [31, 47, 11], many solely rely on evolutionary relationships [16, 24, 6, 10] and are the focus of the current study.

One of the most important distinctions in evolutionary relationships among genes is between orthologs and paralogs [19]. Orthologous genes originate via a speciation event, while paralogous genes arise through a duplication event. By definition, orthologs are always found in different species, though paralogs can be found in either the same or different species (e.g., when duplication precedes a speciation event; Supplementary Figure S1). Identifying orthologous genes across species is important for many tasks, including the inference of species relationships (but see [14, 33, 56]). In the context of function prediction, orthologs have traditionally taken a privileged role based on the belief that they are more functionally similar to one another than are paralogs [18, 13, 51, 20]. Indeed, prediction of function often proceeds by first identifying orthologs, and sometimes only single-copy orthologs, discarding all genes with other relationships (e.g., [53]).

The idea that orthologous genes share greater functional similarity than do paralogous genes has been termed the “ortholog conjecture” [40]. Historically, this conjecture has rarely been questioned in either evolutionary biology (but see [29, 50, 21]) or function prediction studies (but see [16, 36, 38]). In a previous study, we tested the ortholog conjecture using experimental evidence gleaned from the Gene Ontology (GO) database and gene expression data from 25 different tissues in human and mouse [40]. We found no evidence that orthologs were more functionally similar than paralogs of equivalent levels of protein divergence, and in fact showed that the highest functional similarity was shared by within-species paralogs. A simple model was proposed to explain these data: functional differences evolve over time, so that pairs of genes that have been diverged for a smaller amount of time are more functionally similar, and that genes found in the same species share a cellular context, making their functions more similar [40]. Because within-species paralogs can have a common ancestor much more recently than any particular speciation event (which will delimit the age of orthologs)—and are obviously found within the same species—such genes will share higher levels of functional similarity.

Results questioning the ortholog conjecture have been received with a range of reactions. Some researchers were “baffled” that such an obvious assumption would be tested [48], while others pointed out important limitations in using GO similarity for answering questions about the evolution of function [52, 2, 5]. In the rush to cement the primacy of orthologs, several analyses were reported in which orthologs appeared significantly more similar than paralogs [2, 5, 43, 1, 30]. While orthologs may be more similar than paralogs for some traits (cf. [49]) and for some types of genes [1], the higher similarity of orthologs in some newer datasets was accompanied by either no change in functional similarity over time [2, 5] or the *increase* in functional similarity over time [43]. As a decrease in structural [41, 37] and functional [23, 35, 34, 4, 9, 32] similarity with divergence is a widely expected and observed pattern for both paralogs and orthologs, the patterns of evolution in these studies are indeed baffling. Further examination of several of these studies has uncovered problems with the analyses such that there is either no longer support for the ortholog conjecture [15] or that there was no statistical support for the ortholog conjecture in the first place, as in the case of human-mouse comparisons in Ref. [2].

Further testing of the ortholog conjecture using experimental data must deal with several issues, mostly concerned with the non-random nature of experiments done by individual researchers and in individual species [40, 52, 2]. An important distinction that can help to overcome these issues can be drawn between two different interpretations of the ortholog conjecture, one evolutionary and one predictive. Evolutionarily, questions about the tempo and mode of functional evolution require that the same traits and experimental methods be used in all species considered. If this is not done, then differences in the types of traits studied can bias results in favor of within-species paralogs [40, 52]. For instance, if tails are only studied in mice, then obviously more genes in mice will be predictive of tail-related functions. But a second interpretation of the ortholog conjecture is only concerned with the prediction of function, regardless of the evolutionary history of these functions. For this interpretation of the ortholog conjecture there is no bias associated with the non-random collection of experimental data, as long as we assume that all types of annotations are equally accurate on average. Following from the example given above, if more genes in mice are useful for predicting tail-specific functions, then these genes should be preferred when predicting function.

Regardless of the validity of the ortholog conjecture, its importance to the task of function prediction remains unclear. That is, even if orthologs are slightly better for predicting functions than paralogs, this should not imply that paralogs should be ignored, and *vice versa*. In fact, including paralogs in prediction results in higher accuracy than orthologs alone (e.g., [46]) and methods that include both orthologs and paralogs [16, 17] are some of the most successful in the Critical Assessment of Functional Annotation (CAFA) challenge [42, 28, 57]. Therefore, quantifying the increase in the number and accuracy of functional predictions made possible by including paralogs represents an equally valuable task in the context of the ortholog conjecture. Given a large enough benefit of inclusion, it may be that there is no need to distinguish between orthologs and at least some types of paralogs in the first place.

In this paper we revisit the ortholog conjecture and related questions using experimentally verified functional annotations from almost 43,000 genes. We attempt to control for some of the factors that can bias evolutionary tests of the ortholog conjecture, finding that within-species paralogs are overwhelmingly favored in comparisons between two mammalian species, *Homo sapiens* and *Mus musculus*, and between two yeast species, *Saccharomyces cerevisiae* and *Schizosaccharomyces pombe*. We also quantify the enormous gain in number and accuracy of predictions that is manifested when all types of homologs, and not just orthologs, are included. Our results reaffirm the lack of support for the ortholog conjecture, and further suggest that its accuracy is irrelevant to the task of predicting function.

## 2 Methods

### 2.1 Sequence data and evolutionary relationships

We collected protein-coding genes from *Homo sapiens* (Hs), *Mus musculus* (Mm), *Saccharomyces cerevisiae* (Sc) and *Schizosaccharomyces pombe* (Sp). Ensembl Biomart (release 91, December 2017) and Ensembl Fungimart (release 38, January 2018) gene trees were used to specify different homologous relationships for human-mouse and cerevisiae-pombe comparisons, respectively [55]. A total of 8,606 gene trees contained human and mouse genes, whereas 3,059 gene trees contained cerevisiae and pombe genes.

Homologous relationships between proteins were divided into four main categories, two for homologs found in different species (orthologs and between-species outparalogs) and two for homologs found in the same species (inparalogs and within-species outparalogs). Orthologs were further classified as one-to-one, one-to-many and many-to-many. We used duplication events inferred from gene trees to distinguish between inparalogs and within-species outparalogs: if the duplication event occurred more recently than the speciation event, the protein pairs were identified as inparalogs; otherwise, they were identified as within-species outparalogs (Figure S1). All genes included in the final dataset had at least one type of homologous relationship with another gene. This dataset was composed of 19,514 human genes, 21,398 mouse genes, 4,205 cerevisiae genes and 3,487 pombe genes.

### 2.2 Function data

We used Biological Process (BP), Molecular Function (MF) and Cellular Component (CC) ontologies released by the Gene Ontology (GO) consortium in March 2018 [3, 8]. Functional annotations were obtained from the UniProt-GOA database (release 176, March 2018) [26]. Only annotations supported with evidence codes EXP, IDA, IMP, IPI, IGI, IEP, TAS, or IC were considered. The comparative genomics data obtained from Ensembl uses Ensembl gene IDs, whereas protein functional annotation data obtained from the UniProt-GOA database uses UniProt accession numbers. In some cases, especially for human and mouse, when there were one-to-many mappings from Ensembl gene to UniProt accession numbers, we only kept annotations for the protein with the longest sequence. This gave 22,280 human proteins, 12,859 mouse proteins, 5,135 cerevisiae proteins and 2,669 pombe proteins annotated with at least one functional term from any ontology. A total of 81,332 unique publications were used to assign experimental annotations to these proteins.

Our analysis requires pairs of homologs that both have functional annotations in order to quantify their similarity. Therefore, our final dataset consisted of 8,637 ortholog pairs, 1,917 inparalog pairs, 33,741 between-species outparalog pairs, and 40,021 within-species outparalog pairs for human-mouse comparisons. For cerevisiae and pombe, there were 1,724 ortholog pairs, 2,072 inparalog pairs, 193 between-species outparalog pairs, and 892 within-species outparalog pairs.

We also identified proteins that had been annotated in the same publication or by the same researchers. To do so, we retrieved PubMed identifiers for each protein-term assignment and, ultimately, associated a list of authors with each protein. This list was then used to narrow the analysis only to pairs of proteins that were studied by non-overlapping groups of investigators.

### 2.3 Similarity calculations

#### 2.3.1 Sequence identity

Pairs of protein sequences were aligned using the Needleman-Wunsch algorithm [39], with the BLOSUM62 matrix [25], a gap opening penalty of 11, and a gap extension penalty of 1. Sequence identity was obtained by dividing the number of matches in the alignment by the length of the longer protein sequence.

#### 2.3.2 Functional similarity

To calculate functional similarity between pairs of annotated proteins we employed two groups of similarity measures. The first group uses topological measures of the structure of the GO graph to measure similarities of its subgraphs, whereas the second group uses information-theoretic (probabilistic) measures that further incorporate the database of all characterized proteins with their respective annotations. Each measure considered in this work returns values between [0, 1], with 1 indicating identical annotations. Since our main functional similarity measure is Yang-Clark similarity, we introduce it below. The two alternative measures, i.e., the Maryland bridge similarity [22] and Schlicker’s similarity [44], are defined in Supplementary Materials.

##### Yang-Clark semantic similarity

The Yang-Clark distance metric is based on the previously introduced concept of information content of a subgraph within an ontology [7]. We used the normalized version of this semantic distance, as proposed by Yang et al. [54], to calculate functional similarities between protein pairs. This model first assumes that protein annotations are generated by a probabilistic model; i.e., a Bayesian network that has the same structure as GO, with each node in the ontology treated as a binary random variable. The marginal probability for a consistent subgraph *T* associated with a protein is then defined as

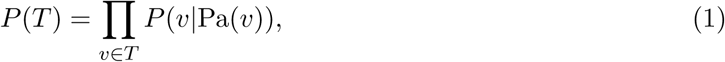

where *v* is a node in the annotation graph and Pa(*v*) is the the set of parent nodes of *v. P* (*v*|Pa(*v*)) defines the probability that a node *v* belongs to the functional annotation of a protein in the database given that all of its parents are present in the annotation. The information content of a subgraph *T* annotating a protein is then defined as

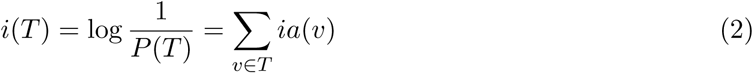

where *ia*(*v*) = − log *P* (*v*|Pa(*v*)) stands for information accretion of a node [7]. We estimated *ia*(*v*) using the maximum likelihood approach over the entire UniProt-GOA database as the negative binary logarithm of the relative frequency that the term *v* is present in a protein’s annotation given that all its parent terms are also present.

The semantic distance between annotations of two proteins can now be calculated as follows. Suppose that *A* and *B* are two sets of nodes presenting propagated annotations for two proteins *a* and *b*, respectively, and that *A* is used as a prediction of *B. Misinformation* (*mi*) is defined as the total information content of the nodes that are present in annotation *A* but not in *B*, whereas *remaining uncertainty* (*ru*) is defined as the total information content of the nodes that are present in annotation *B* but not in *A* [7]. More formally,

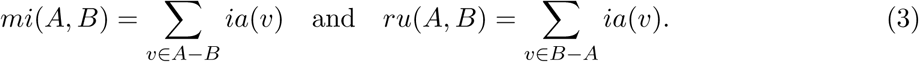

We define the total normalized Yang-Clark distance of the order *p ≥* 1 between the two annotations *A* and *B* as

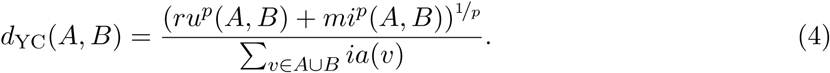

The Yang-Clark semantic similarity is then defined as

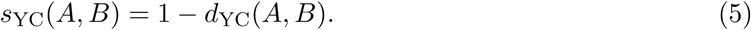

Based on previous work [7, 27, 54], we used *p* = 2 throughout this study.

#### 2.3.3 Background similarity

Background functional similarity is defined as the expected functional similarity for a pair of randomly selected genes. We calculate such similarity for different groups of genes with the same labels, such as orthologs or inparalogs. To calculate background similarity we randomly selected with replacement 1000 protein pairs from a pool of all proteins forming a certain group; e.g., the pool of all proteins forming human-mouse orthologous pairs. The average functional similarity over these pairs can then be subtracted from the functional similarity of the actual homologous pairs to form the so-called *excess similarity* [2].

We calculated background similarities separately for each homology type. Orthologs were further split between one-to-one and other orthologs (one-to-many, many-to-many). Background similarities for non-homologous genes both within and between species were also calculated.

### 2.4 Protein function prediction and its evaluation

To understand and quantify the influence of particular types of homologs in protein function prediction, we selected one of the most intuitive predictors used in this field. This predictor transfers protein annotations from a database of experimentally annotated “target” proteins to an unannotated “query” protein as follows: (1) the query protein is first aligned to all target proteins; (2) each annotation term is transferred from the target database to the query with a score equaling the largest global sequence identity between the query and any of the target proteins containing that term. Eventually, each term in the query protein is associated with a score between 0 and 1. This model is equivalent to the BLAST baseline algorithm that has been used in the CAFA experiments [42, 28, 57], except that we used global sequence identity instead of local identity. We selected this algorithm because of the ability to easily track the influence of target proteins from which the annotations were transferred.

The performance of protein function prediction was evaluated using a leave-one-out strategy based on both topological and information-theoretic accuracy measures. As in pairwise functional similarity between proteins, the Yang-Clark semantic similarity was used as the main evaluation metric and is presented below. The *F*_max_ topological metric, a criterion regularly seen in CAFA [42, 28, 57], was used as an alternative measure and is presented in Supplementary Materials.

To summarize this performance evaluation, we consider a prediction algorithm on a set of *n* proteins, where each protein *i* is assigned a score (say, between 0 and 1) for each functional term *v* in the ontology. The normalized remaining uncertainty (*nru*) and misinformation (*nmi*) are defined as

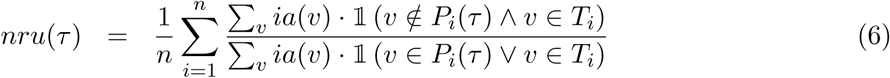

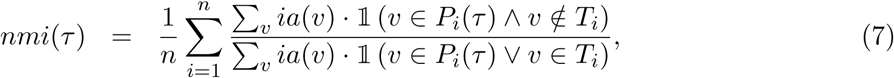

where *P*_*i*_(*τ*) contains predicted terms with a score greater than or equal to *τ* for the *i*-th protein, *T*_*i*_ is the experimental annotation for the *i*-th protein, and 𝟙 is an indicator function. The term *ia*(*v*), as before, is the information accretion corresponding to the ontology term *v* [7]. The maximum semantic similarity, *S*_max_, is now defined as

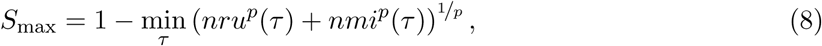

for *p ≥* 1. *S*_max_ takes values between 0 and 1. Higher values correspond to better predictions, with the value of 1 corresponding to a perfect prediction for each protein in the data set. As before, we used *p* = 2.

It is important to mention that all functional similarity measures between proteins are susceptible to problems caused by incomplete [12] and noisy [45] experimental annotations. There is a small effect of annotation incompleteness on topological measures and a somewhat larger effect on unnormalized semantic distance [27]. However, compared with topological measures, semantic similarity avoids a form of double-counting of nodes caused by the directed acyclic graph structure of GO, and thus properly treats hierarchical dependencies in the ontology.

## 3 Results

### 3.1 Higher functional similarity in within-species homologs

We analyzed patterns of functional similarity vs. sequence identity for pairs of proteins separated by their type of homology: orthologs, inparalogs, within-species outparalogs, and between-species outparalogs (Figures 1 and 2). We observe that for both pairs of species and across all three functional ontologies, within-species homologs—especially inparalogs—generally exhibit higher average functional similarity than between-species homologs (orthologs and between-species outparalogs). Inparalogs by definition share a more recent common ancestor with each other than do orthologs, and it therefore may not be surprising that they are more functionally similar than pairs of orthologs at essentially all levels of divergence (cf. [40]). Outparalogs are not constrained by such relationships, but our results show that within-species pairs are consistently more functionally similar than are between-species pairs. This result is also consistent with the previously proposed effect of cellular and organismal context on measured protein function [40]. The patterns, shown for the Yang-Clark similarity measure in Figures 1 and 2, also hold when using Maryland bridge and Schlicker’s similarity measures (Figures S2 and S3).

**Figure 1:**
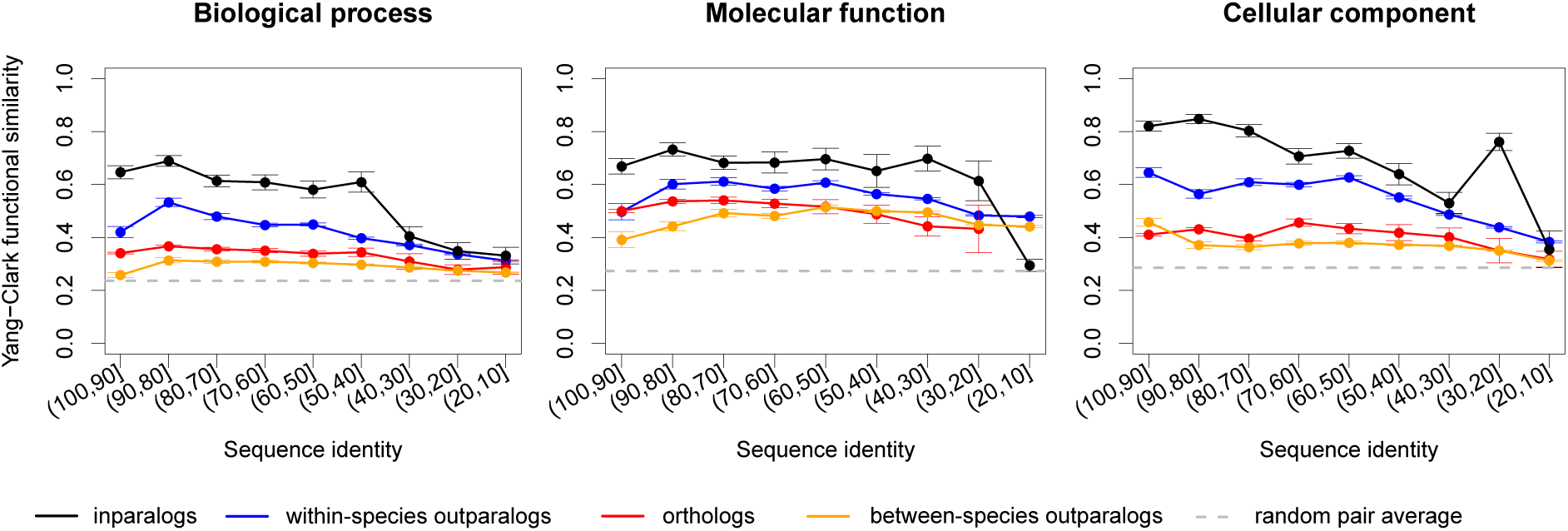
The relationship between Yang-Clark functional similarity and sequence identity for human and mouse over three different ontologies in GO. The breakdown is presented for four types of homologous relationships between pairs of proteins (orthologs, between-species outparalogs, within-species outparalogs, and inparalogs). The dashed line in each panel shows the estimated functional similarity for a randomly selected pair of proteins, obtained by averaging 1000 randomly selected proteins from the available pool. The data are presented for each sequence identity bin in which at least three pairs of proteins could be used to calculate functional similarity.

**Figure 2:**
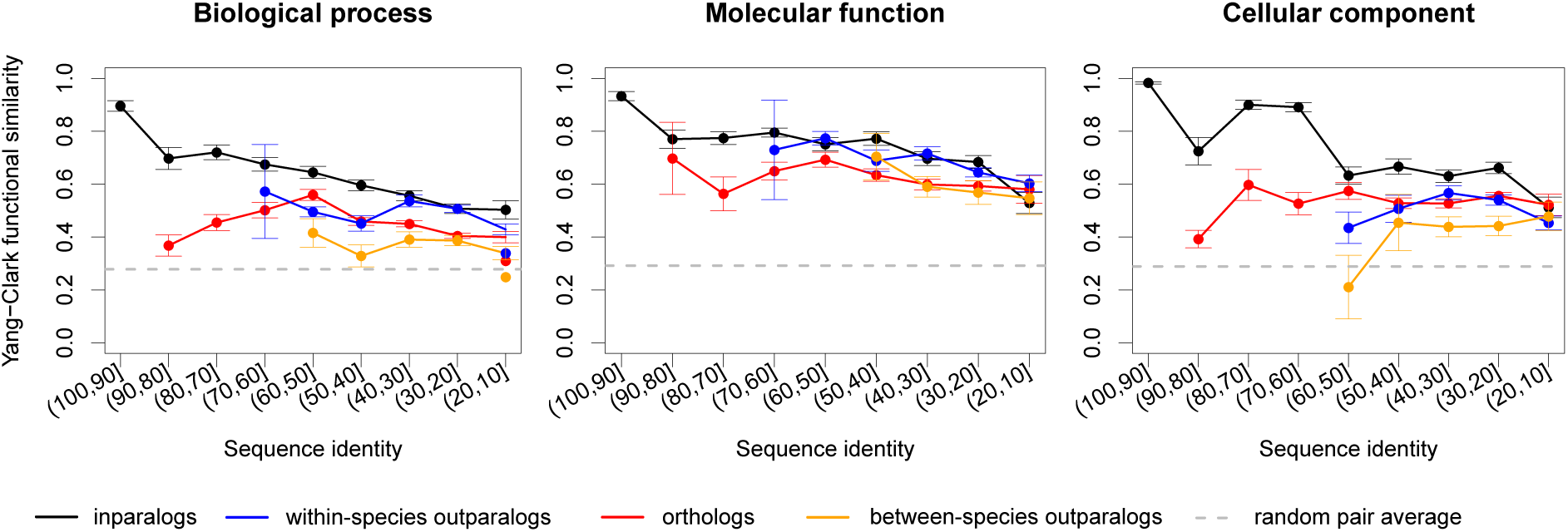
The relationship between Yang-Clark functional similarity and sequence identity for *S. cerevisiae* and *S. pombe* over three different ontologies in GO. The breakdown is presented for four types of homologous relationships between pairs of proteins (orthologs, between-species outparalogs, within-species outparalogs, and inparalogs). The dashed line in each panel shows the estimated functional similarity for a randomly selected pair of proteins, obtained by averaging 1000 randomly selected proteins from the available pool. The data are presented for each sequence identity bin in which at least three pairs of proteins could be used to calculate functional similarity.

### 3.2 Controlling for potential annotation bias

While the results presented in Figures 1 and 2 clearly show that within-species homologs have more annotated functional terms in common than do between-species homologs, it is not clear whether this is due to underlying biological differences. We studied two sources of bias that could inflate functional similarities of within-species homologs, in an attempt to control for them. The first factor examined was “authorship bias,” which proposes that pairs of proteins experimentally annotated by the same authors will be annotated more similarly [40, 2]. This effect could be due either to a limited range of GO terms known by individual authors or to experiments preferentially testing similar functions. In either case proteins studied by the same authors could have higher functional similarity compared to pairs annotated by different authors. The second potential factor is referred to as “background similarity” between pairs of homologs [2]. Such similarity arises because certain functions are studied more or less in different organisms, leading to similarities even between non-homologous proteins found in the same species.

To examine the effects of these factors on our results, we first calculated functional similarity between pairs of proteins annotated by different authors. We did this by removing any homologous pair for which experimental annotations were derived from either the same paper or different papers sharing one or more of their authors (see Methods). Across all ontologies, and for both species pairs, functional similarity for inparalogs remained higher than for orthologs for high sequence similarity, whereas the results were mixed in the lower sequence identity groups (Figure S4). The effect of filtering of functional annotations was more noticeable for within-species outparalogs: these become comparable in functional similarity to orthologs across the entire sequence identity range, with fluctuations likely influenced by smaller dataset sizes (Figure S4). Finally, between-species outparalogs remained the least predictive of functional annotations (Figure S4). These results were consistent regardless of the similarity measure used (Figures S4-S6). To control for background similarity, we calculated this measure separately for the different types of homologous relationships (Figures S7-S9). We then subtracted the background similarity from the total functional similarity between homologous pairs of the same type; these values were subtracted from the similarities calculated above using annotations only from different authors. Inparalog pairs remained functionally slightly more similar to each other than orthologs after accounting for both sources of biases (using all three measures and over all three ontologies; Figures S10-S12). While there remain several unavoidable problems with experimental data collected from different species (see Discussion), these results suggest that orthologs do not evolve functions more slowly than paralogs.

### 3.3 The impact of ignoring paralogs: number of genes

Regardless of whether paralogs are more or less functionally similar than orthologs of the same sequence identity, using only orthologs for function prediction means that we are discarding some amount of functional information. We therefore attempted to characterize the impact of discarding paralogs both in terms of the number of proteins that are ignored and the loss of predictive ability. As before, these results are shown for human vs. mouse and cerevisiae vs. pombe. Using Ensembl gene trees [55], we extracted 8,606 gene families for human-mouse and 3,059 for cerevisiae-pombe that contained at least two proteins within them of some homology type. We were interested in quantifying the number of families and the number of proteins in the two pairs of species where functional transfer is possible given: (i) only orthologs, (ii) only paralogs, and (iii) both orthologs and paralogs.

Figure 3 shows a significant overlap between protein families where functional transfer is possible using both orthologs and any type of paralogs. In human and mouse, a total of 4,744 families with 9,488 proteins contain only orthologous assignments, whereas 260 families with 1,396 proteins contain only paralogous assignments, mostly from the same species (i.e., either human-human or mouse-mouse). On the other hand, 3,602 gene families containing 30,000 proteins contain both types of homologs and thus could potentially benefit from both groups in function transfer. For *S. cerevisiae* vs. *S. pombe*, we find that 1,969 gene families (3,968 proteins) only have orthologous assignments, 571 families (1,590 proteins) only have paralogous assignments, whereas 519 families (1,772 proteins) contain both types of homologous assignments. Additional breakdowns are provided in Supplementary Materials.

**Figure 3:**
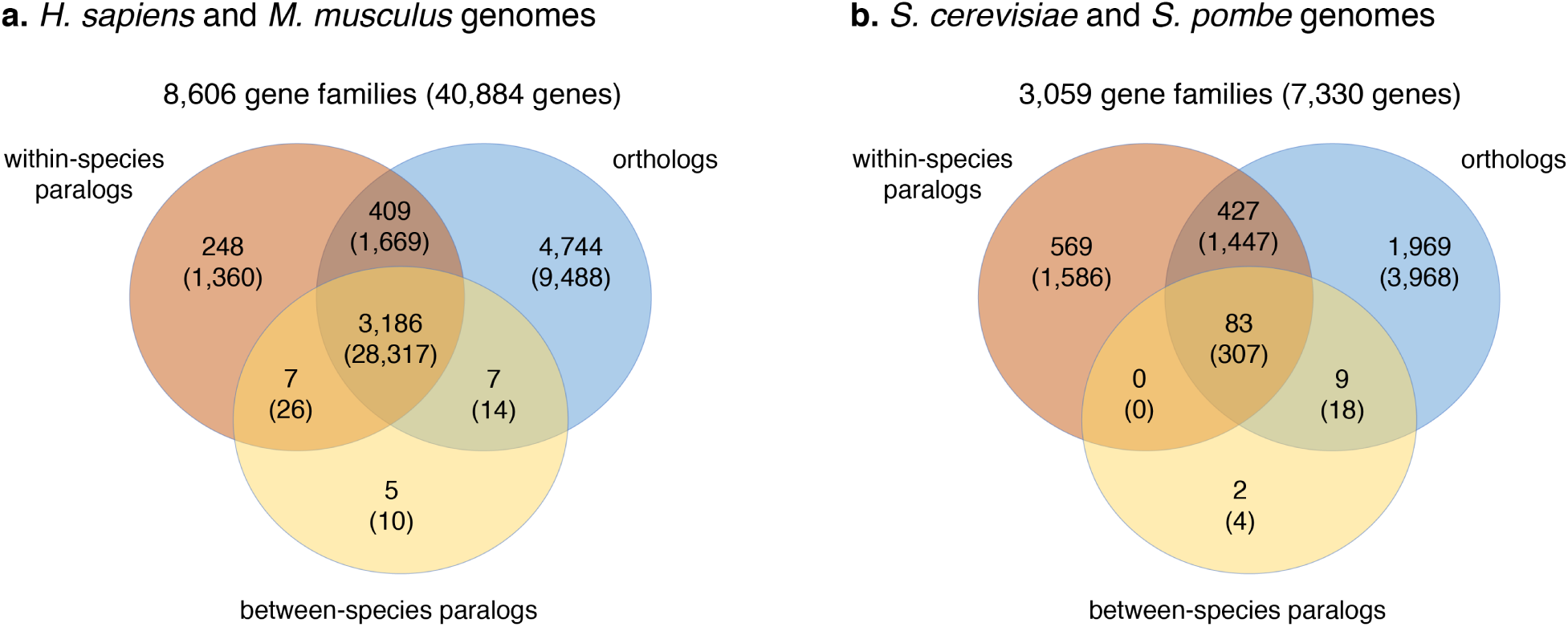
The numbers of gene families with different types of homologous relationships in them. The numbers in parentheses represent the counts of genes in the respective gene families. (a) gene families containing *H. sapiens* and *M. musculus* proteins, and (b) gene families containing *S. cerevisiae* and *S. pombe* proteins.

These results point to the potentially large impact paralogs could have on function transfer. Although orthologs are the only source of function transfer in 55% of human and mouse families and 64% of yeast families, 73% of proteins in human and mouse and 24% of proteins in the yeast species are in families containing genes with both orthologous and paralogous relationships. In addition, 3% of families in human and mouse (3% of proteins) contain only paralogous relationships and 19% of families (22% of proteins) in the yeast species contain only paralogous proteins. Together, these numbers suggest that a large amount of functional information is potentially ignored if paralogs are not included in function transfer.

### 3.4 Impact of ignoring paralogs: prediction performance

We estimated the performance accuracy of protein function prediction when different groups of targets (i.e., different types of homologs) were used to transfer GO terms. We first estimated the prediction accuracy when orthologs and all paralogs were used for the prediction. We then removed orthologs and paralogs one at a time to gauge the impact on prediction performance. Figure 4a-c shows the prediction performance in each ontology for the proteins that have an experimentally characterized ortholog and at least one paralog, regardless of the type (gray circles). Removing orthologs (blue circles) resulted in significantly reduced performance across all ontologies and across both pairs of species (10.2% reduction on average in human-mouse and 13.4% reduction on average in cerevisiae-pombe). Similarly, removing paralogs from function transfer (yellow circles) resulted in reduced performance by an equivalent margin to that for the case of orthologs (10.0% reduction on average in human-mouse and 16.8% on average in cerevisiae-pombe). These results suggest that the two groups of homologs provide approximately equivalent contributions to accurate function transfer.

**Figure 4:**
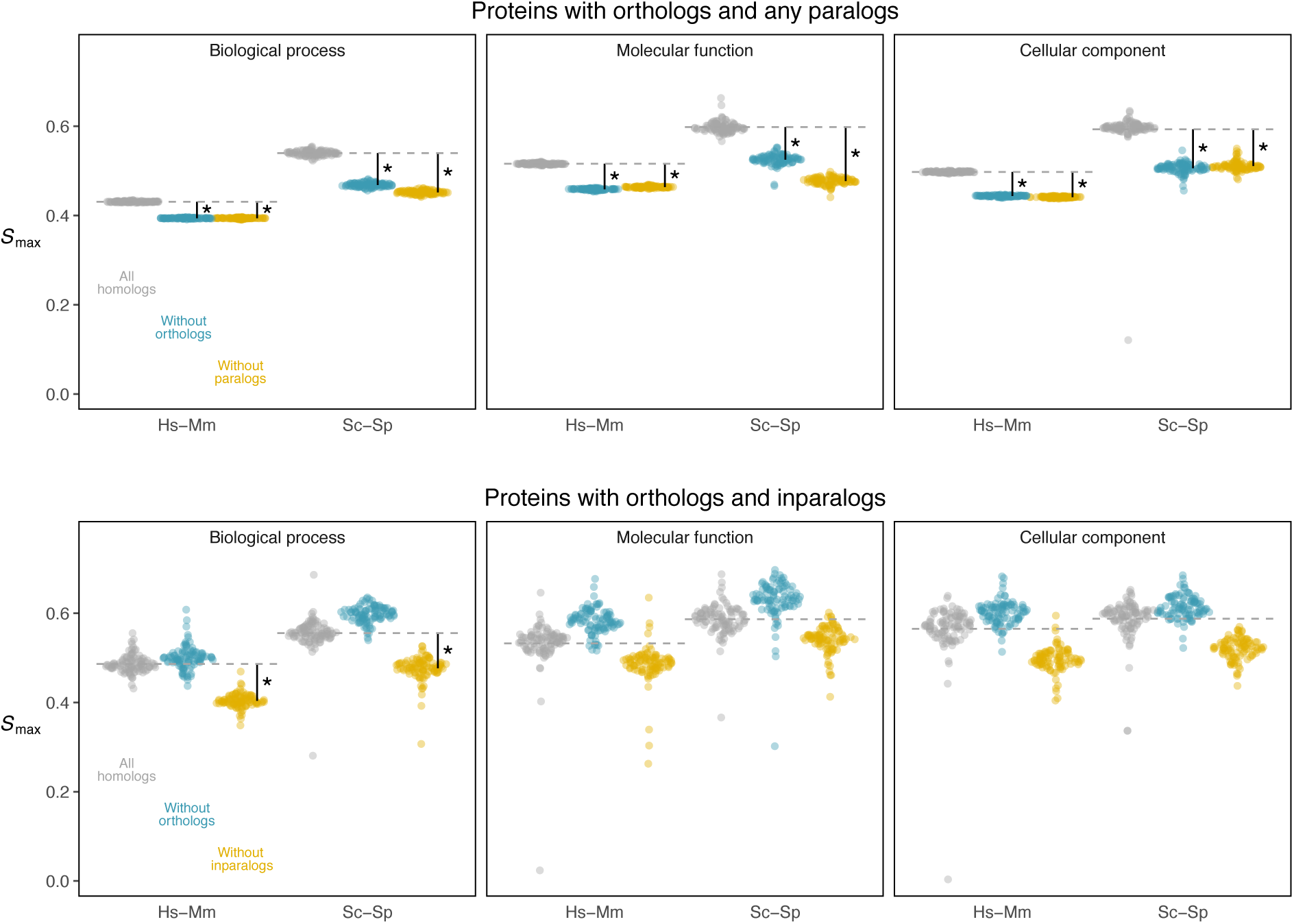
The impact of excluding orthologs or paralogs from prediction. *S*_max_ values for proteins having orthologs and one other homolog (top panels) and proteins having orthologs and inparalogs (bottom panels), in human-mouse (Hs-Mm) and *S. cerevisiae*-*S. pombe* (Sc-Sp), are lower when ignoring either homolog. Points are 100 bootstrap samples of each protein set, and asterisks indicate a significant decrease in *S*_max_ (bootstrap *P <* 0.05).

Figure 4d-f summarizes a similar experiment, but where we only considered proteins with both orthologs and inparalogs having experimentally determined functions. Once again, the removal of inparalogs resulted in a significant decrease in prediction performance across all ontologies and both pairs of species (12.6% reduction on average in human-mouse and 11.1% reduction on average in cerevisiae-pombe). Interestingly, however, the exclusion of orthologs from function transfer resulted in slight performance increases in all six experiments, although this result was not statistically significantly (6.2% increase on average in human-mouse and 6.5% increase on average in cerevisiae-pombe). These results suggest that the annotated inparalogs, when available, constitute the most reliable source of functional annotation and can be readily used in function transfer. The results show that accuracy actually decreased when orthologs were included alongside the inparalogs. Further experiments with the removal of other types of homologs and different performance measures are summarized in the Supplementary Materials (Figures S13-S24).

## 4 Discussion

We have revisited the problem of the “ortholog conjecture,” with a focus on assessing the value of orthologs and paralogs in the task of sequence-based protein function prediction. Exploiting significantly larger data sets of experimentally characterized proteins than have been used before, we repeated the analysis of [40], finding that those original results still hold for two different pairs of species. We then moved beyond this analysis to quantify the value of different types of homologs on both the number of possible predictions that can be made and their accuracy. Several conclusions and implications of our results deserve additional discussion.

We believe that the ortholog conjecture as originally operationalized in our previous work conflated two different ideas about the value of orthologs: their role in automated function prediction versus their rate of evolution of function [40]. Although these are obviously related issues, different types of experiments—and more importantly, different types of experimental biases—can affect the conclusions drawn about the roles of orthologs in each. Questions about the role of orthologs in functional annotation have been satisfactorily answered, with no problems due to experimental bias: the cumulative evidence suggests that paralogous genes are highly important for functional transfer, and in certain cases even more useful than orthologs in transferring function from one gene and/or species to another. The analyses presented here continue to show that paralogs can offer both more and better predictions of protein function; that is, given a query protein and experimentally annotated orthologs and paralogs, the combination of orthologs and paralogs consistently improves functional annotation compared to prediction based on orthologs alone. These results hold across both a comparison of human and mouse and a comparison of two yeast species. We [40] and others [2] have previously shown the same patterns in human and mouse, though Ref. [2] did not find the same pattern in their analysis of the two yeasts. We do not know exactly why the two results are different, but our work is based on six more years of accumulated data.

In contrast, questions regarding the evolution of function in different types of homologs are still unsettled. The results presented here are consistent with a model in which protein functions evolve over time, with no difference between orthologs and paralogs except with respect to whether two genes are found within the same species [40]. In previous work there was also no evidence for a greater conservation of function in orthologs between human and mouse when using GO terms, even after correcting for experimental evidence coming from the same paper [40] or from the same authors [2]. However, there remain issues present in the experimental data reported in the GO database that are hard to overcome in downstream analyses of protein function evolution. One such issue is that current measures of functional similarity treat a lack of overlap in GO terms as evidence for a difference in function, without properly accounting for the possibility that the relevant experiments have simply not been carried out [52]. Such problems can only be addressed as more experiments are added to the GO database, especially experiments reporting negative results. Indeed, by tracking the accumulation of data in GO, Chen and Zhang [5] predicted that the functional similarity of ortholog pairs would be higher than for outparalog pairs by the year 2018 for all three ontologies. We used experimental data from more than six times as many publications in this study than in our previous study [40] and did not find evidence for this (Figure 1 and Figure 2). Moreover, Chen and Zhang [5] predicted that functional similarity between orthologs would exceed those of inparalogs for the biological process and cellular context ontologies in 2013 and 2015, respectively; these predictions are also not supported by our results. Nevertheless, further work is clearly necessary to directly assess the rate of evolution of protein function among different classes of homologs.

The results presented here suggest an important modification to approaches for assigning protein function. Regardless of the validity of the ortholog conjecture for this task—that is, no matter whether orthologs are better or worse than paralogs at predicting function—adherence to the ortholog conjecture is often accompanied by the idea that only orthologs should be used to predict function [48]. This is clearly not a necessary condition of the ortholog conjecture, as orthologs could have more conserved function while still including paralogs in function prediction tasks. As we have shown, ignoring paralogs involves throwing away a huge amount of data, and often means that no prediction can be made because there are no available orthologs (Figure 3). Stricter schemes can also involve only using one-to-one orthologs (Figure S25), so as to avoid the inclusion of any duplication events in the history of the orthologs. This strategy means that even fewer genes can be used in function prediction.

As sequence and function data further accumulate, future analyses, particularly those including multiple species at a time, could reveal more refined relationships between types of homology and function. Until such a time, however, we propose that homology types should be ignored in methods for transferring protein function, with a caveat that the functions from between-species outparalogs are slightly less transferable from one species to another. Our results suggest both that we are ignoring large amounts of data and that the accuracy of prediction is lower if we do not use paralogs. Even studies written in favor of the ortholog conjecture provide only slim-at-best victory margins [2, 5, 43, 1, 30], while still presenting data that support the value of paralogs. Furthermore, evolutionary relationships among genes (i.e., a gene tree) can still be used to predict function, even when the labelling of orthologs and paralogs is ignored [17]. Such approaches are some of the most accurate in function prediction [42, 28, 57], and support the idea that having more high-quality data will almost always improve prediction accuracy. While distinguishing between orthologs and paralogs is a necessary step for answering important biological questions, we find both groups to be predictive of protein function and therefore valuable for function transfer.

## Acknowledgements

The authors acknowledge the support by the NSF grant DBI-1458477 and the Precision Health Initiative of Indiana University.

