## Supplementary Materials for "The Ortholog Conjecture Revisited: the Value of Orthologs and Paralogs in Function Prediction"

### Supplementary Methods

This document provides supplementary information for the manuscript "The Ortholog Conjecture Revisited: the Value of Orthologs and Paralogs in Function Prediction" by Moses Stamboulian, Rafael F. Guerrero, Matthew W. Hahn, and Predrag Radivojac.

**Maryland bridge similarity.** Let $A$ and $B$ be two sets of GO terms denoting the propagated functional annotations for the proteins $a$ and $b$, respectively. The Maryland bridge similarity [1] is defined as

$$s_{\mathrm{M}}(A, B) = \frac{1}{2} \cdot \left( \frac{|A \cap B|}{|A|} + \frac{|A \cap B|}{|B|} \right), \tag{1}$$

and can be interpreted as an average of precision and recall when the terms associated with protein $a$ are used as a prediction of the terms associated with protein $b$, and *vice versa.* The root term was excluded from these calculations, in order to exploit the full $[0, 1]$ range for functional similarity.

**Schlicker semantic similarity.** The Schlicker similarity used here is a slight modification of the original similarity measure [2], as proposed in [3]. To calculate the similarity between two leaf GO terms, $t_i$ and $t_j$, this measure uses Lin's similarity measure defined as

$$s_{\mathrm{L}}(t_i, t_j) = \frac{2 \cdot s_{\mathrm{R}}(t_i, t_j)}{\log {}^1\!/_{P(t_i)} + \log {}^1\!/_{P(t_j)}}, \tag{2}$$

where $P(t)$ is the probability of observing a term $t$ in a randomly selected protein, and $s_{\mathrm{R}}(t_i, t_j)$ is Resnik's similarity [4] between the terms, calculated as

$$s_{\mathrm{R}}(t_i, t_j) = \max_{t \in \mathcal{A}(t_i, t_j)} \frac{1}{\log P(t)}. \tag{3}$$

In Eq. (3), $\mathcal{A}(t_i, t_j)$ indicates the set of common ancestor terms between terms $t_i$ and $t_j$ in the ontology. Term probabilities were estimated using their relative frequencies in the annotations of all species in the UniProt-GOA database.

As gene products are often annotated with more than one leaf term, functional similarity for a protein pair was calculated using the best match average method, proposed originally in Schlicker et al. [2], giving the following Schlicker similarity

$$s_{\mathrm{S}} = \frac{1}{|p_1| + |p_2|} \cdot \left( \sum_{i \in p_1} \max_{j \in p_2} (s_{\mathrm{L}}(i, j)) + \sum_{j \in p_2} \max_{i \in p_1} (s_{\mathrm{L}}(i, j)) \right) \tag{4}$$

Combining the terms in Eq. (4) differs from the original formula suggested in [2] in the way that the two maximum terms are averaged. This particular implementation was chosen to closely match the semantic similarity from [3].

**Topological metrics.** The performance of protein function prediction was evaluated using a leave-one-out strategy based on one topological and one information-theoretic accuracy measure. We used the $F_{\max}$ topological metric, a criterion regularly seen in CAFA [5, 6]. To summarize this performance evaluation process, we consider a prediction algorithm on a set of $n$ proteins, where each protein $i$ is assigned a score (say, between 0 and 1) for each functional term $v$ in the ontology. We now define an average precision ($pr$) and recall ($rc$) at some score threshold $\tau$ as

$$pr(\tau) = \frac{1}{m(\tau)} \sum_{i=1}^{m(\tau)} \frac{\sum_v \mathbb{1}(v \in P_i(\tau) \wedge v \in T_i)}{\sum_v \mathbb{1}(v \in P_i(\tau))} \quad (5)$$

$$rc(\tau) = \frac{1}{n} \sum_{i=1}^{n} \frac{\sum_v \mathbb{1}(v \in P_i(\tau) \wedge v \in T_i)}{\sum_v \mathbb{1}(v \in T_i)}, \quad (6)$$

where $P_i(\tau)$ contains predicted terms with a score greater than or equal to $\tau$ for the $i$-th protein, $T_i$ is the experimental annotation for the $i$-th protein, $m(\tau)$ is the number of proteins having at least one annotation transferred with a score greater than or equal to $\tau$, and $\mathbb{1}$ is an indicator function. The $F_{\max}$ score is defined as

$$F_{\max} = \max_{\tau} \left\{ \frac{2 \cdot pr(\tau) \cdot rc(\tau)}{pr(\tau) + rc(\tau)} \right\} \quad (7)$$

and represents the maximum harmonic mean between average precision and average recall over all thresholds.

### Supplementary Figures

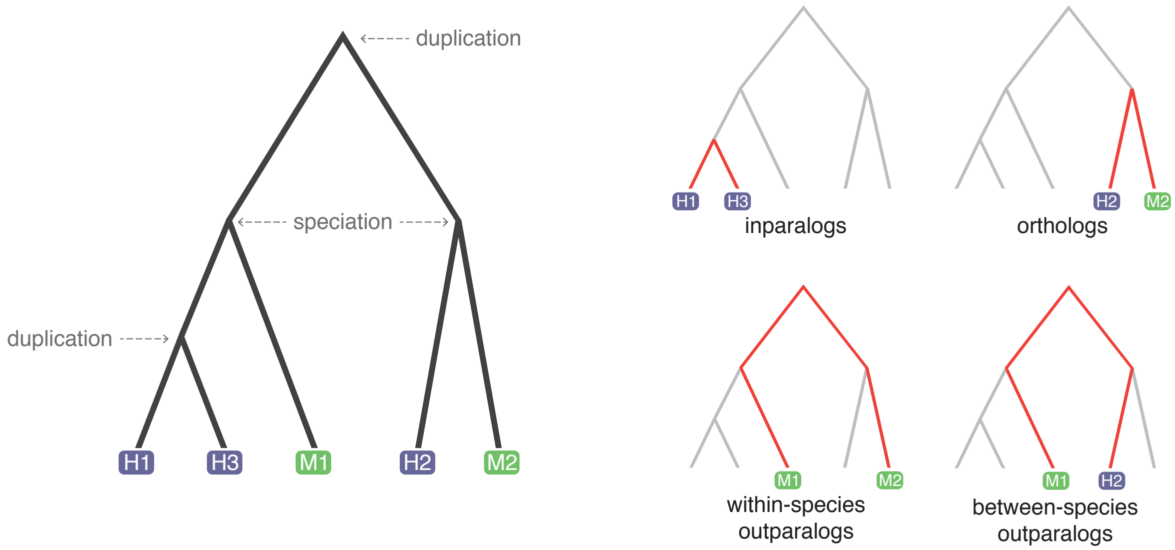


Figure S1: Four different types of homology relations. A family of five genes sampled from human (in blue) and mouse (in green) evolves through speciation and duplication events (left-hand tree). The relationships among genes are highlighted in the four trees on the right. Human genes H1 and H3 are inparalogs, arising from a duplication event after the most recent speciation event in the tree. Genes H2 and M2 are orthologs, as they are related through a speciation event. Gene pairs M1-M2 and M1-H2 are outparalogs because they arose from a duplication that predates the reference speciation event. Pair M1-M2 are within-species outparalogs, while M1-H2 are between-species outparalogs.

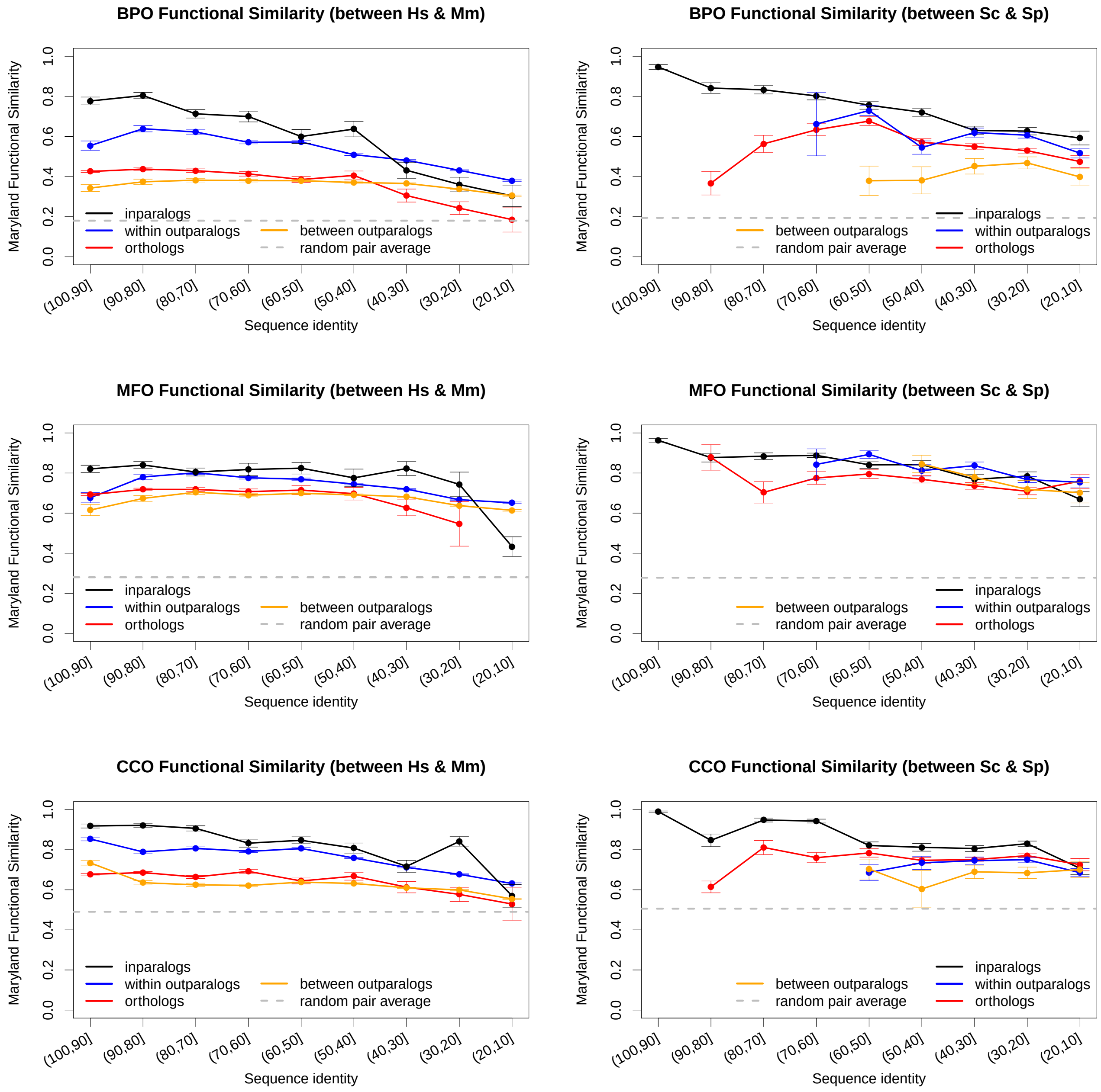


Figure S2: Relationship between functional similarity and sequence identity for human-mouse and cerevisiae-pombe homologs for all three ontologies, using Maryland bridge similarity.

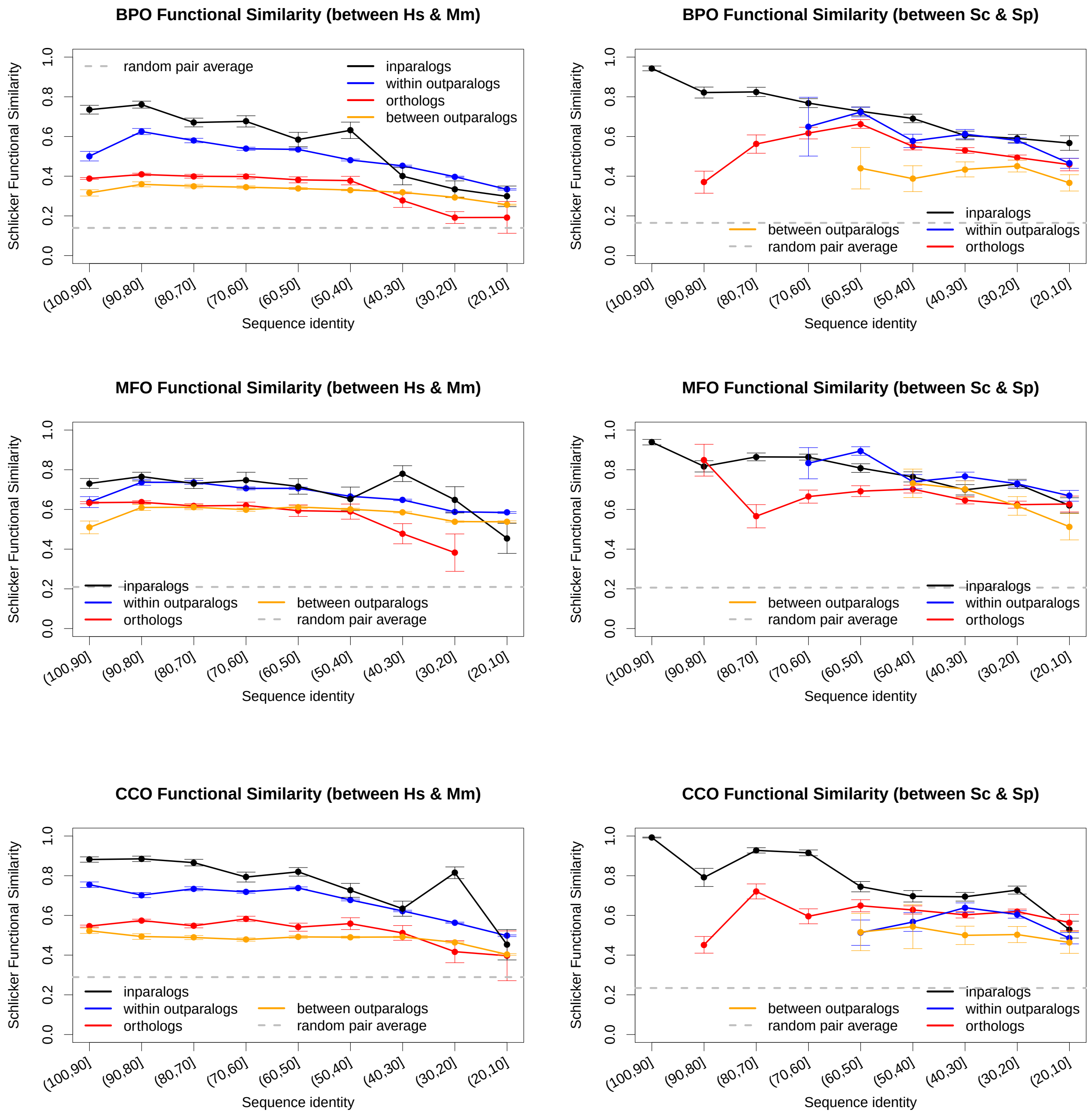


Figure S3: Relationship between functional similarity and sequence identity for human-mouse and cerevisiae-pombe homologs for all three ontologies, using Schlicker's similarity.

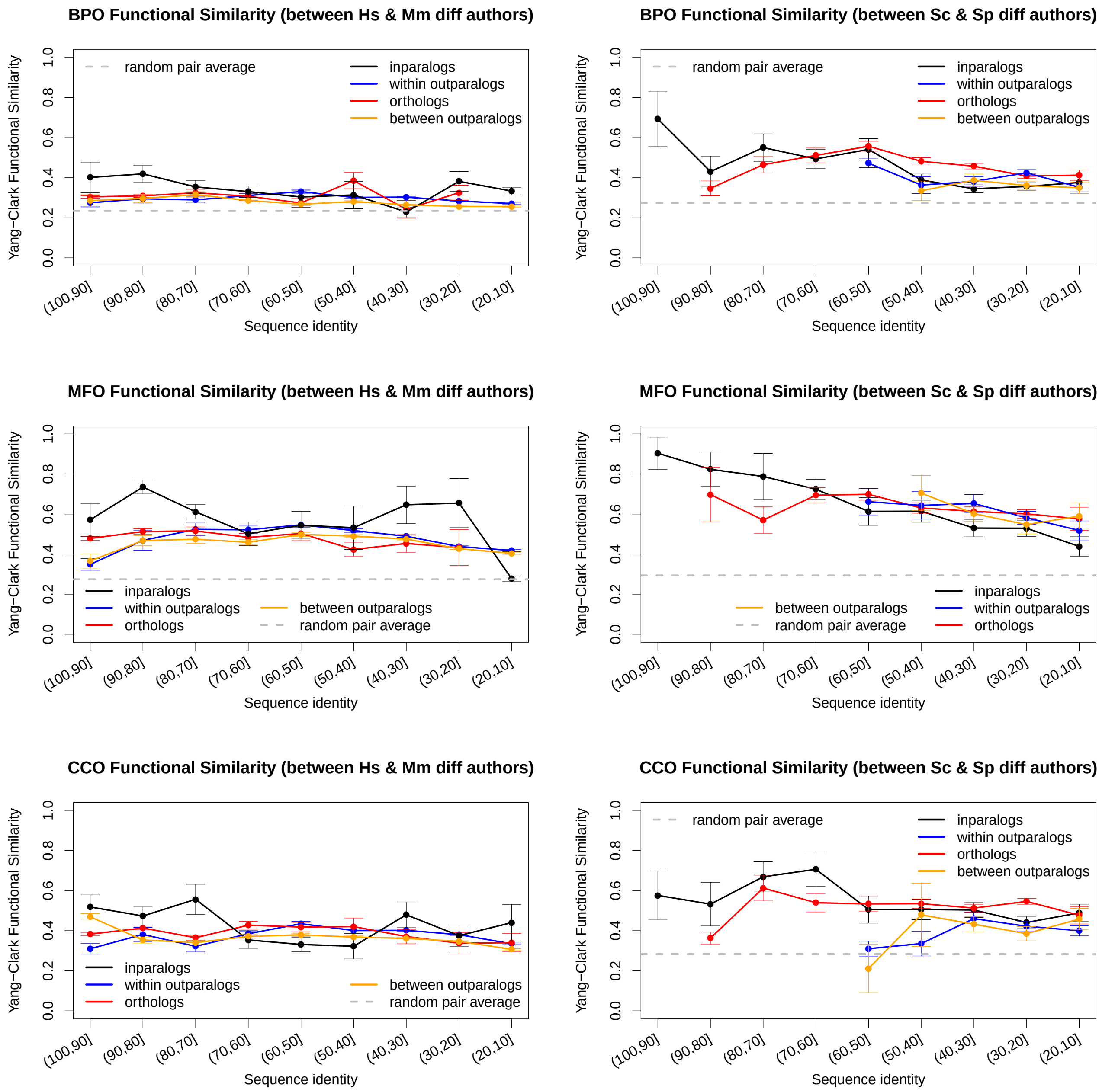


Figure S4: Relationship between functional similarity and sequence identity for human-mouse and cerevisiae-pombe pairs annotated by different authors, for all three ontologies, using Yang-Clark similarity.

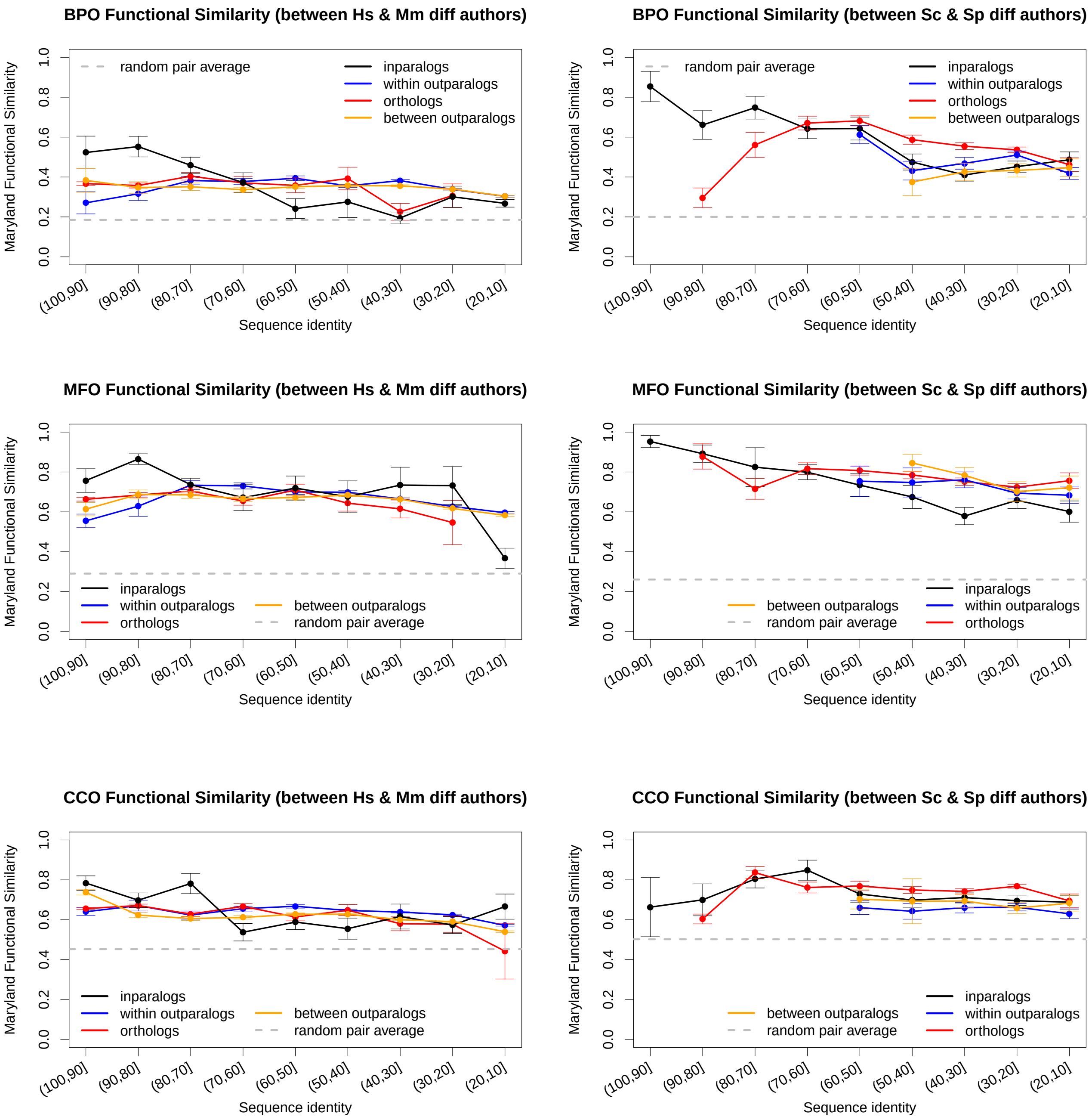


Figure S5: Relationship between functional similarity and sequence identity for human-mouse and cerevisiae-pombe pairs annotated by different authors, for all three ontologies, using Maryland bridge similarity.

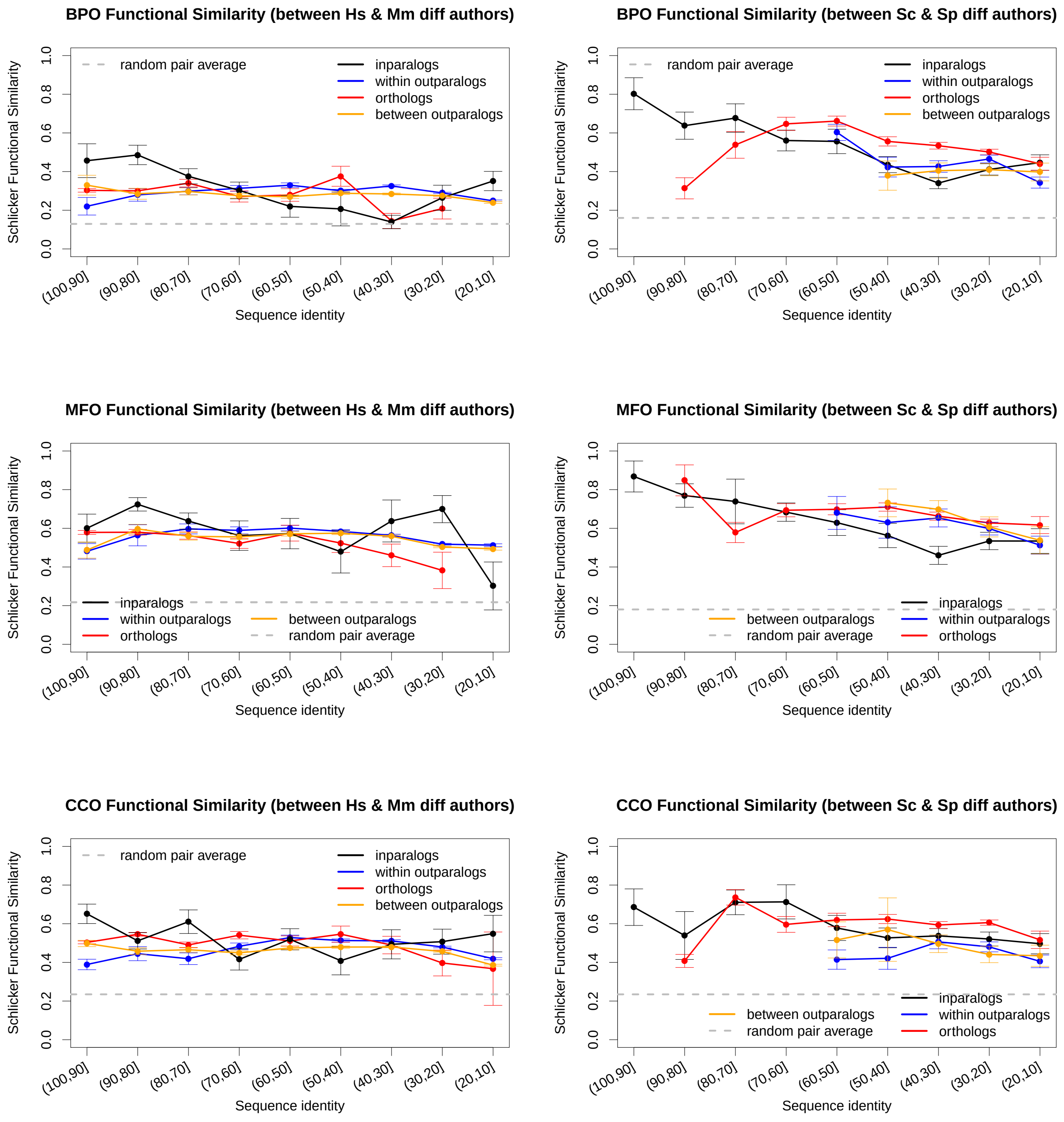


Figure S6: Relationship between functional similarity and sequence identity for human-mouse and cerevisiae-pombe pairs annotated by different authors, for all three ontologies, using Schlicker's similarity.

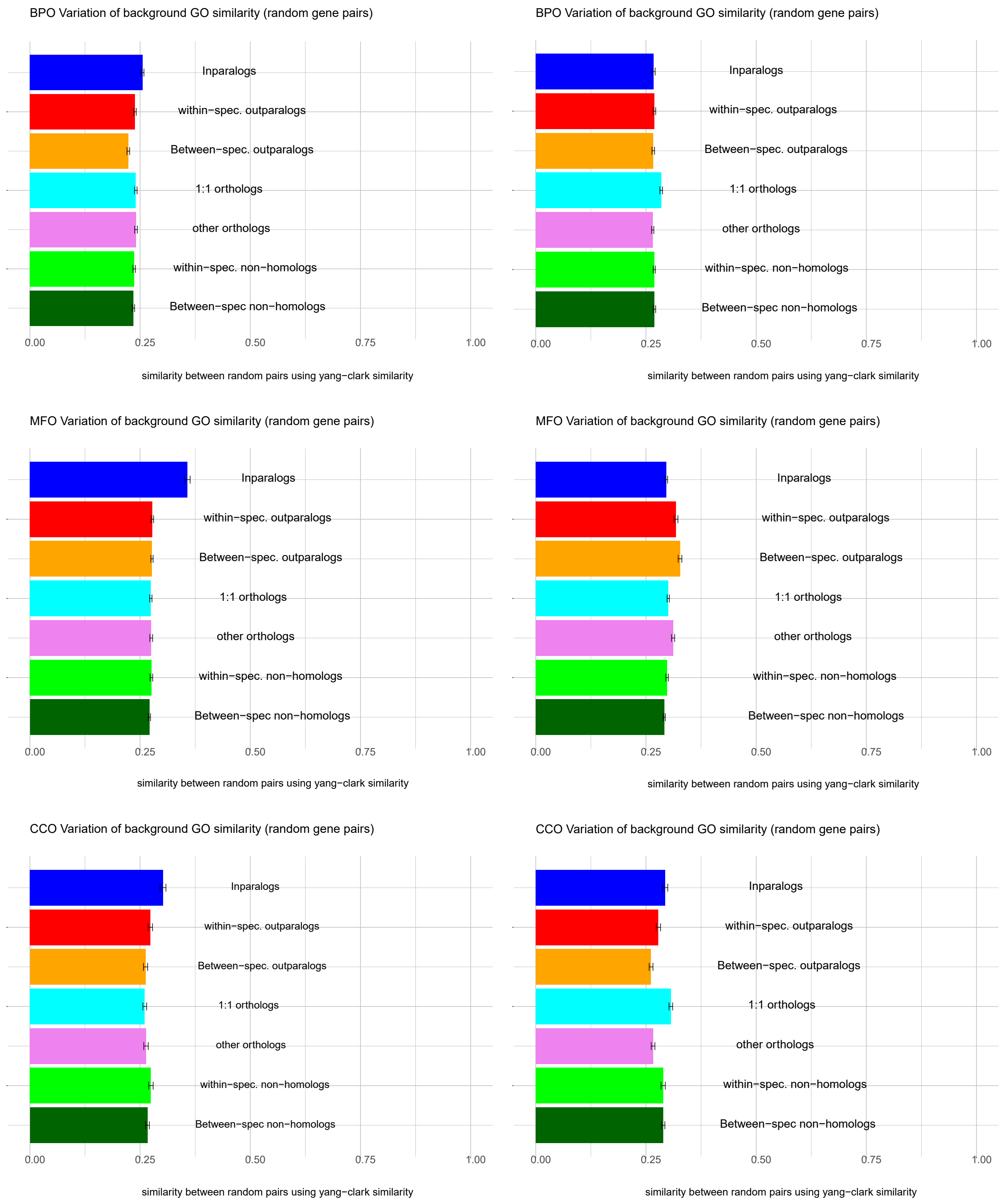


Figure S7: Background similarity for different types of homology relationships for human-mouse (left panels) and cerevisiae-pombe (right panels), using Yang-Clark similarity.

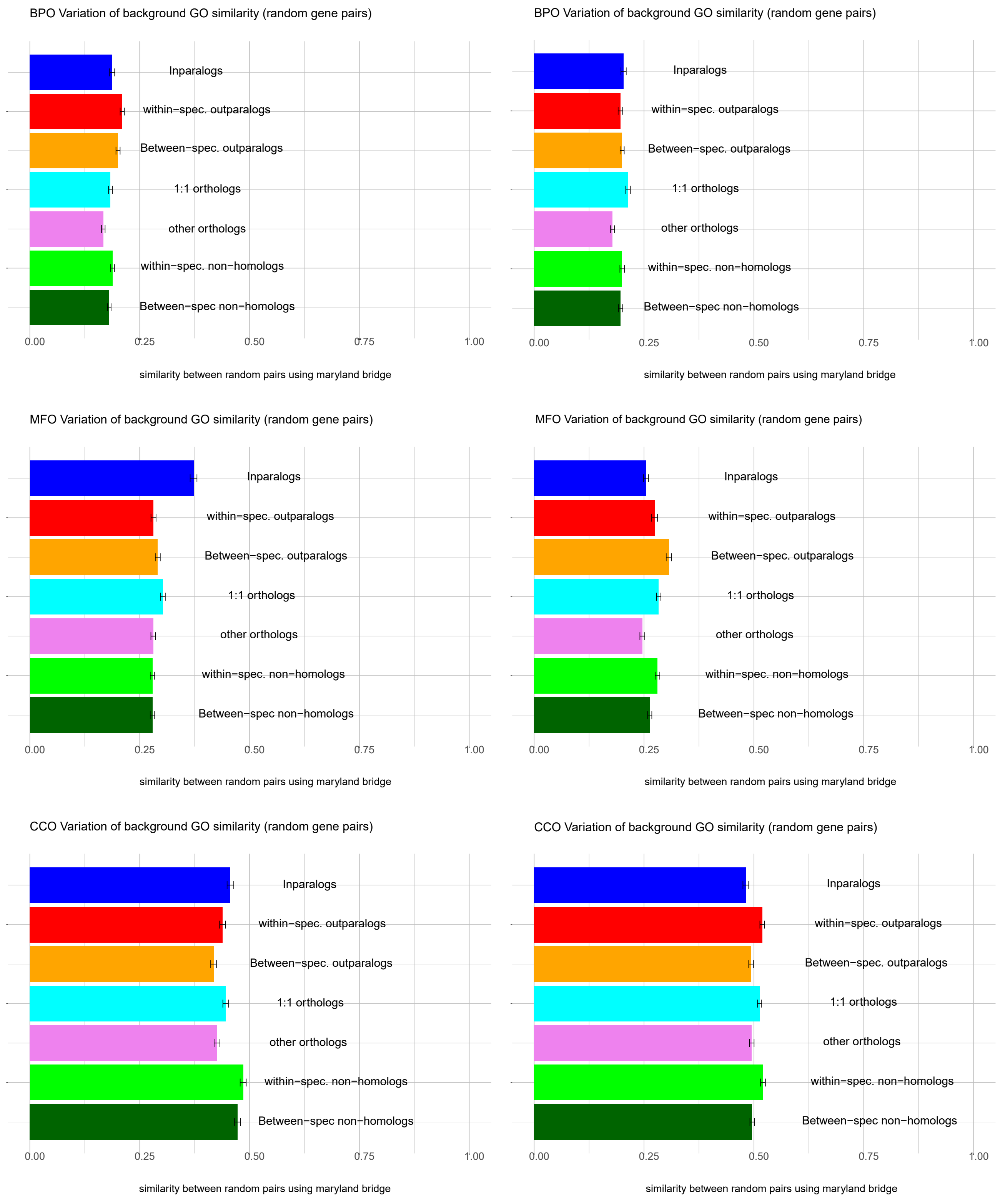


Figure S8: Background similarity for different types of homology relationships for human-mouse (left panels) and cerevisiae-pombe (right panels), using Maryland bridge similarity.

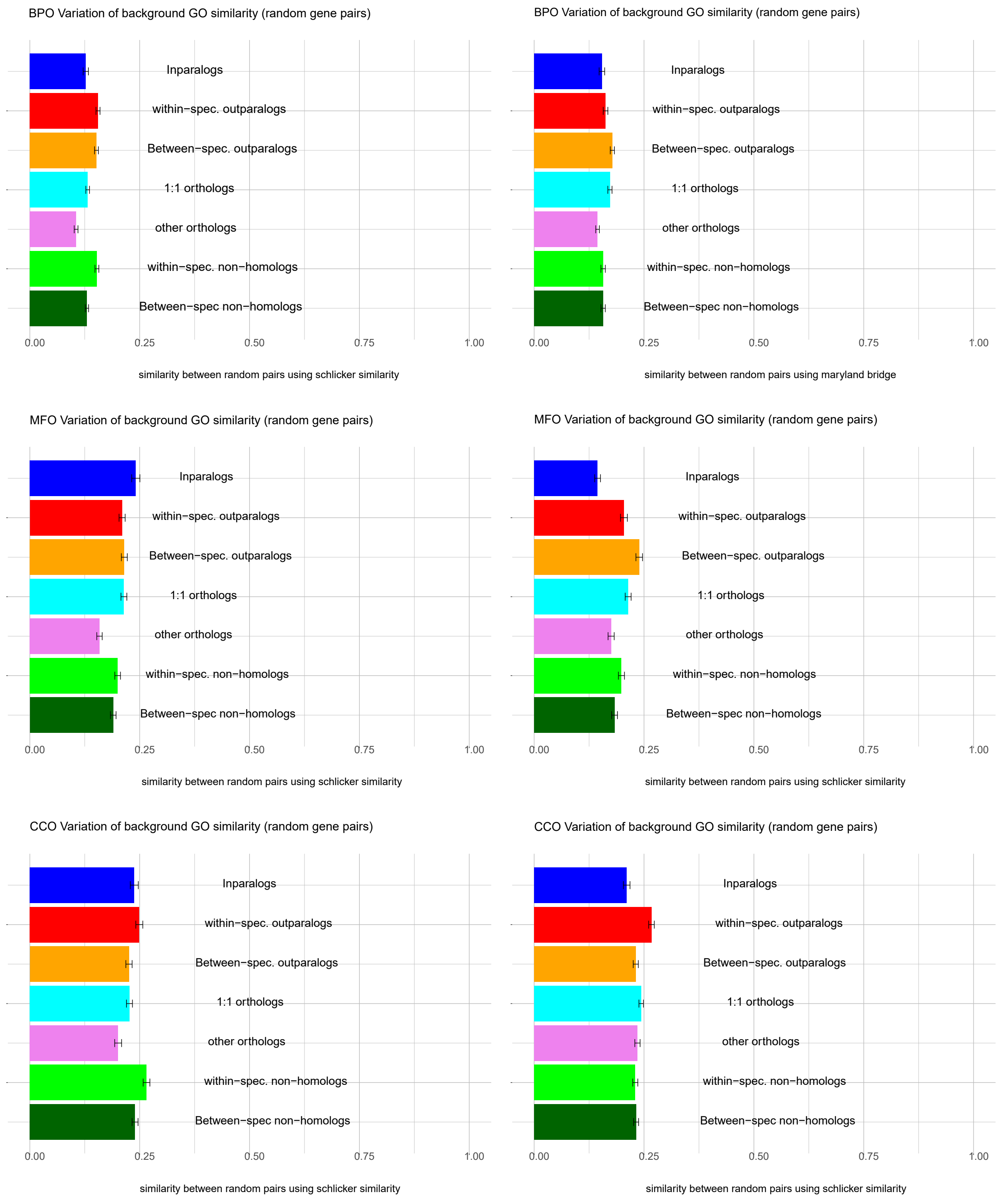


Figure S9: Background similarity for different types of homology relationships for human-mouse (left panels) and cerevisiae-pombe (right panels), using Schlicker's similarity.

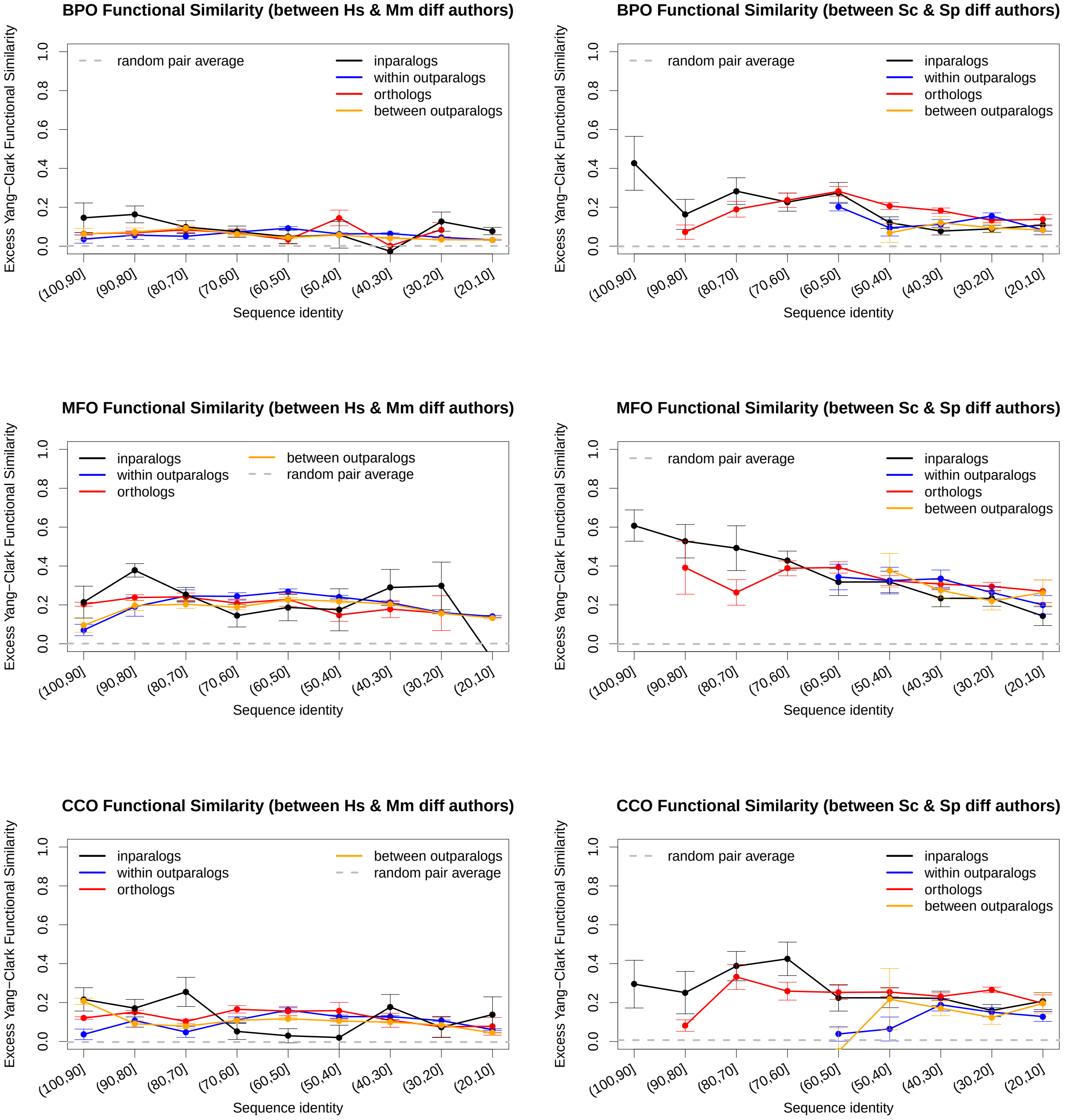


Figure S10: Relationship between functional similarity and sequence identity for human-mouse and cerevisiae-pombe pairs annotated by different authors after removing background similarities, for all three ontologies, using Yang-Clark similarity.

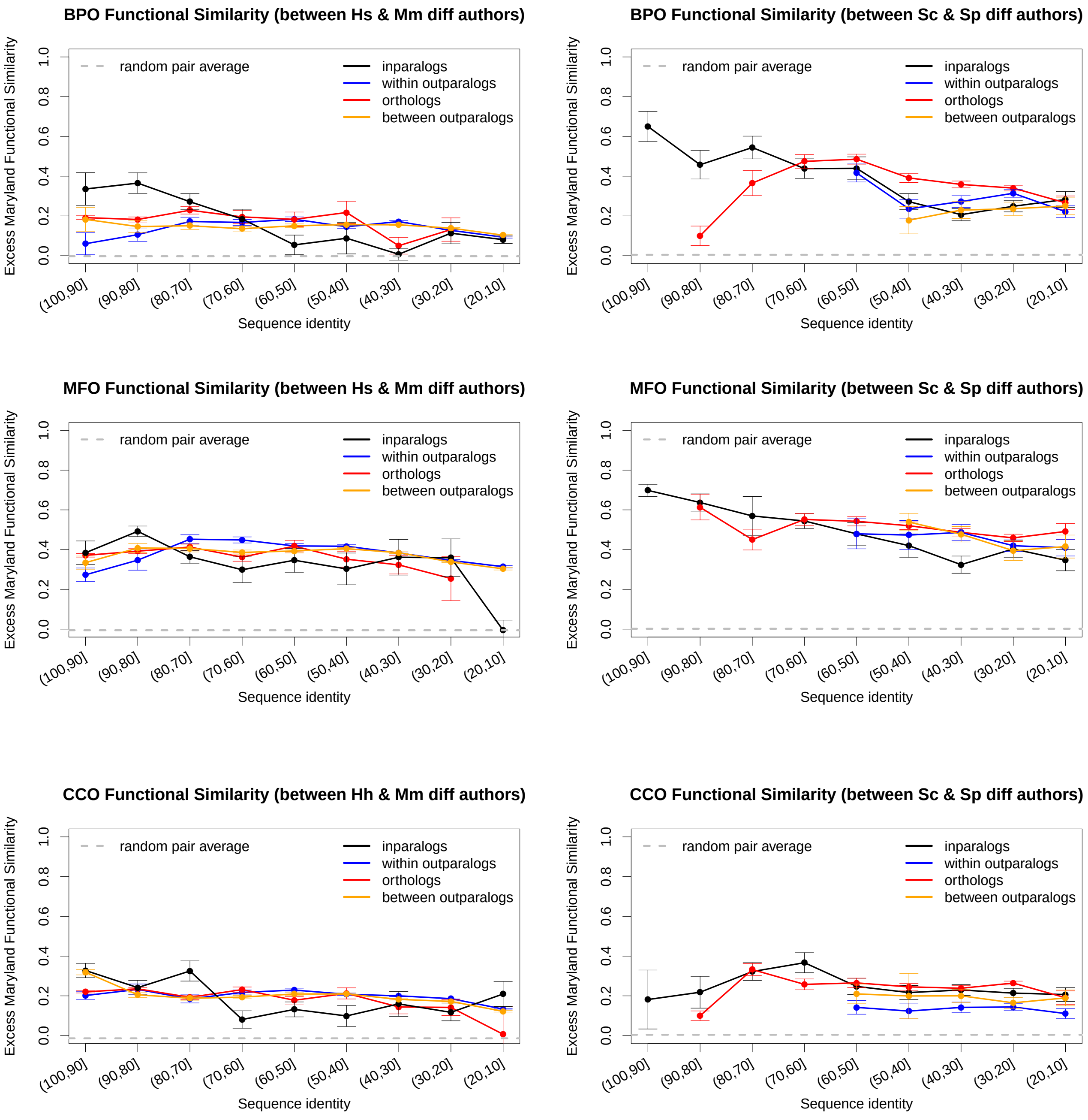


Figure S11: Relationship between functional similarity and sequence identity for human-mouse and cerevisiae-pombe pairs annotated by different authors after removing background similarities, for all three ontologies, using Maryland bridge similarity.

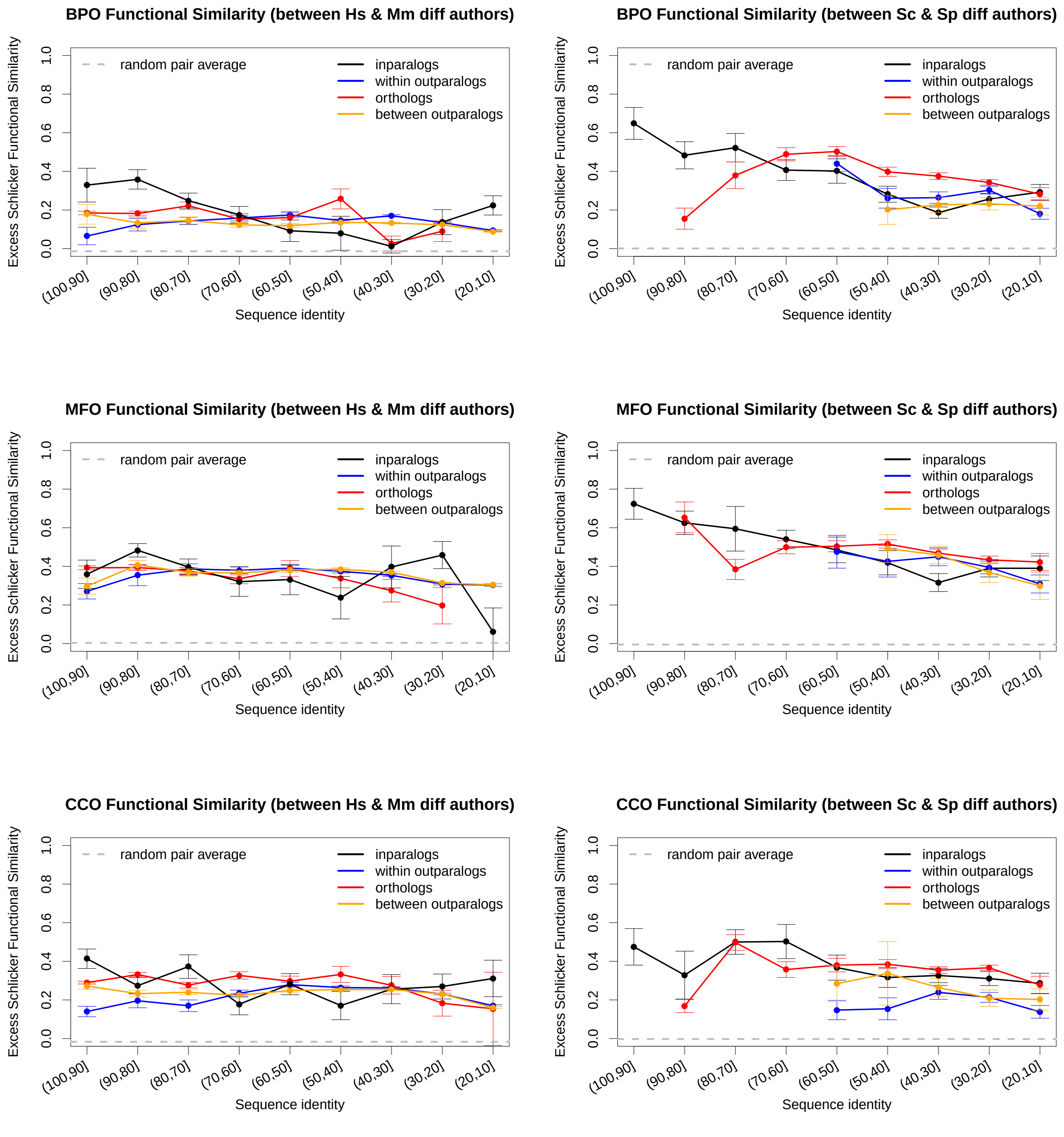


Figure S12: Relationship between functional similarity and sequence identity for human-mouse and cerevisiae-pombe pairs annotated by different authors after removing background similarity, for all three ontologies, using Schlicker's similarity.

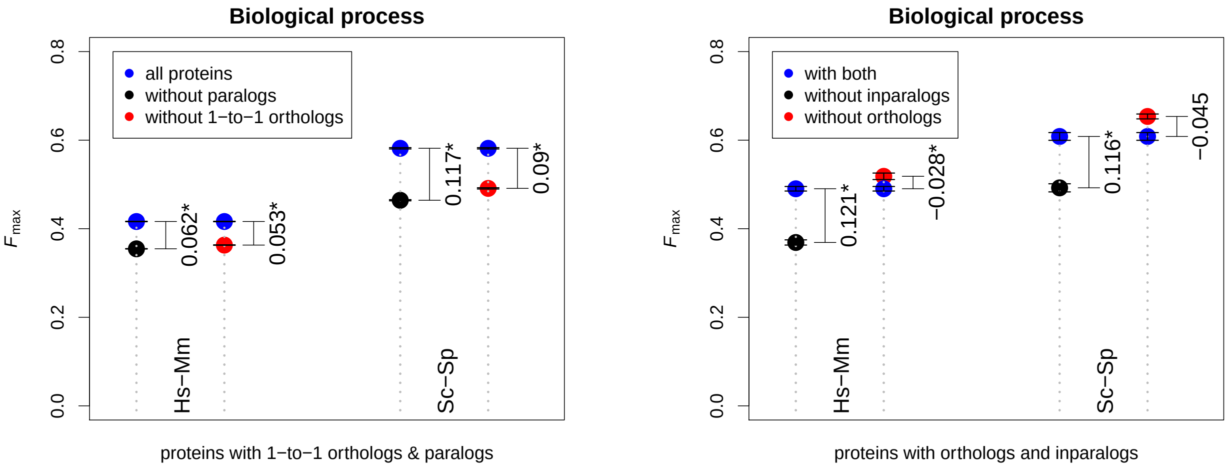


(a) (b)

Figure S13: $F_{\max}$ values for proteins having orthologs, proteins having orthologs and other homologs (a) and proteins having 1-to-1 orthologs and paralogs (b) in human-mouse and cerevisiae-pombe using biological process ontology (* indicates values are statistically significant).

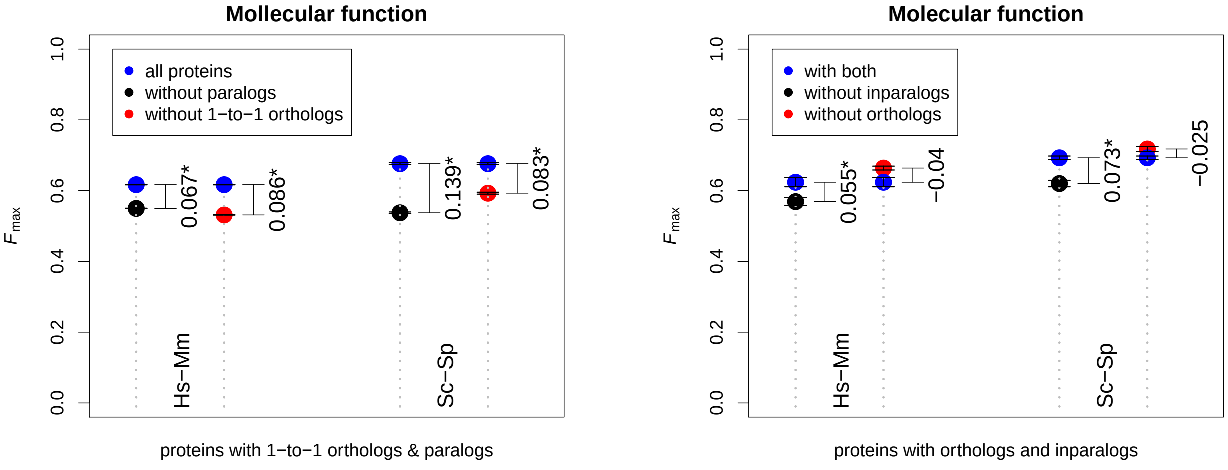


(a) (b)

Figure S14: $F_{\max}$ values for proteins having orthologs, proteins having orthologs and other homologs (a) and proteins having 1-to-1 orthologs and paralogs (b) in human-mouse and cerevisiae-pombe using molecular function ontology (* indicates values are statistically significant).

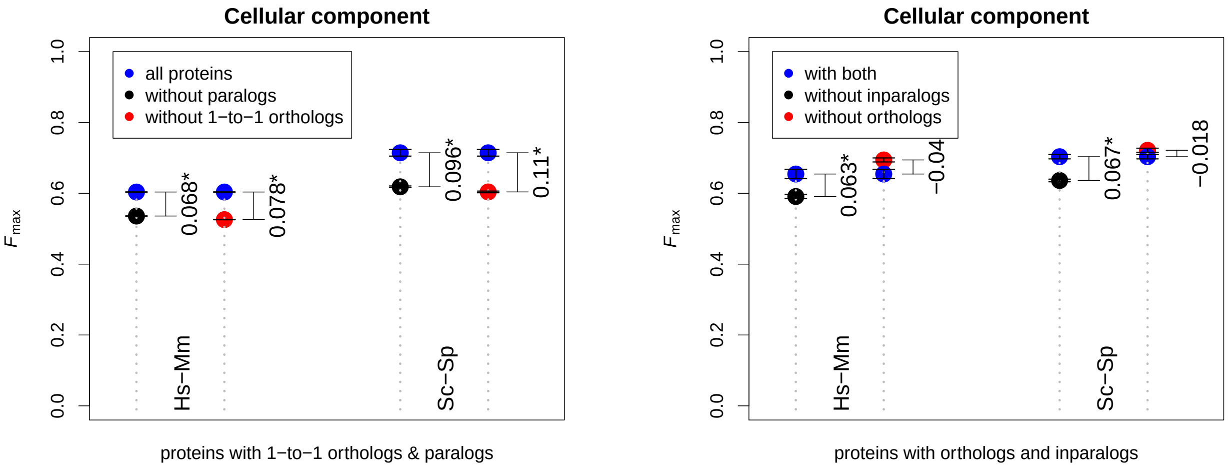


(a) (b)

Figure S15: $F_{\max}$ values for proteins having orthologs, proteins having orthologs and other homologs (a) and proteins having 1-to-1 orthologs and paralogs (b) in human-mouse and cerevisiae-pombe using cellular context ontology (* indicates values are statistically significant).

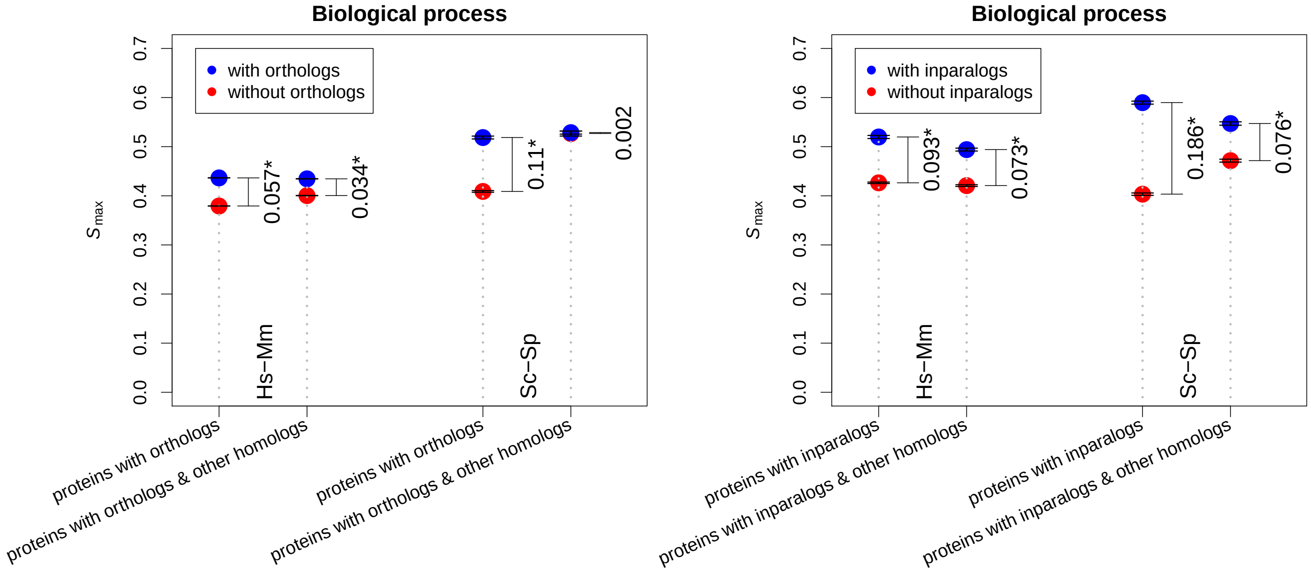


(a) (b)

Figure S16: $S_{\max}$ values for proteins having inparalogs, proteins having inparalogs and other homologs (a), and proteins having both inparalogs and orthologs (b) in human-mouse and cerevisiae-pombe using biological process ontology (* indicates values are statistically significant).

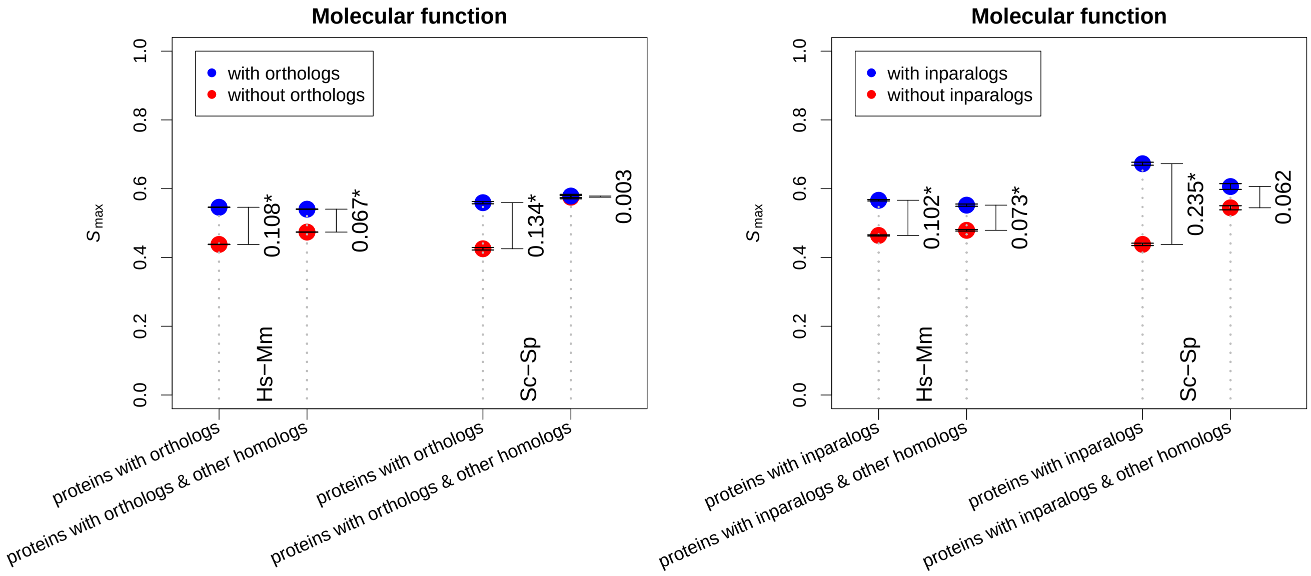


(a) (b)

Figure S17: $S_{\max}$ values for proteins having inparalogs, proteins having inparalogs and other homologs (a), and proteins having both inparalogs and orthologs (b) in human-mouse and cerevisiae-pombe using molecular function ontology (* indicates values are statistically significant).

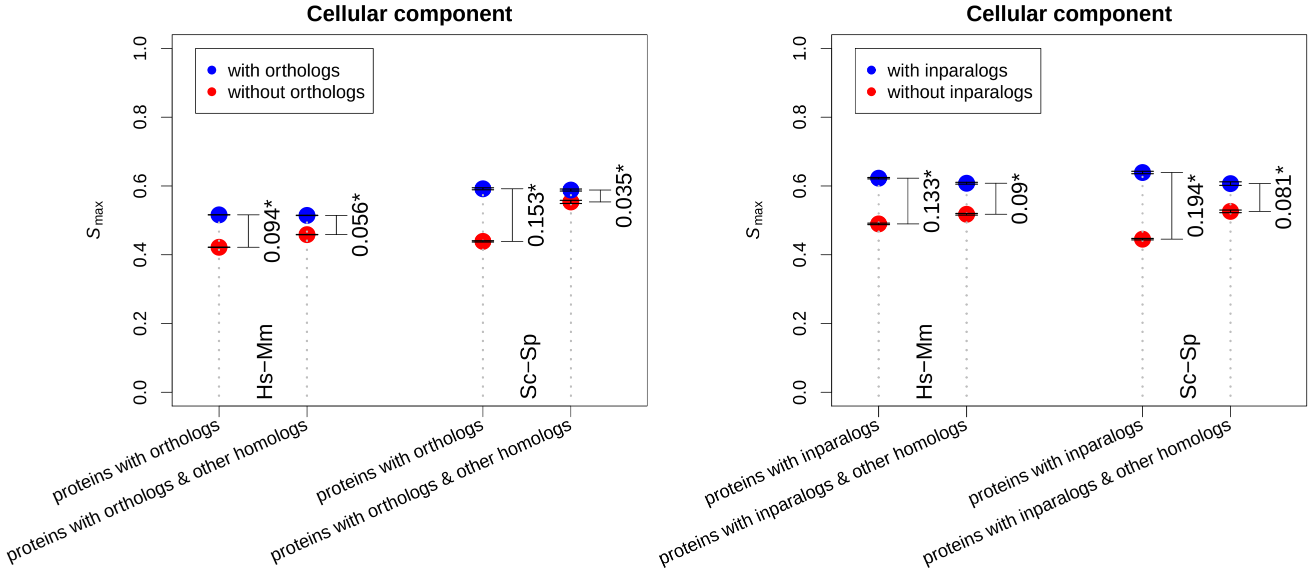


(a) (b)

Figure S18: $S_{\max}$ values for proteins having inparalogs, proteins having inparalogs and other homologs (a), and proteins having both inparalogs and orthologs (b) in human-mouse and cerevisiae-pombe using cellular context ontology (* indicates values are statistically significant).

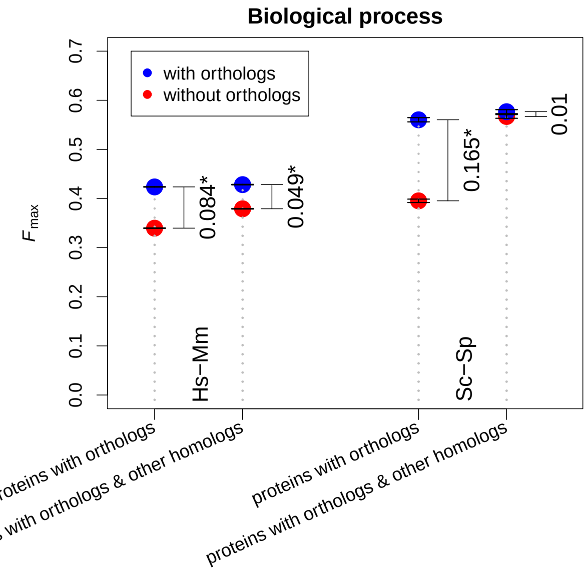


(a)

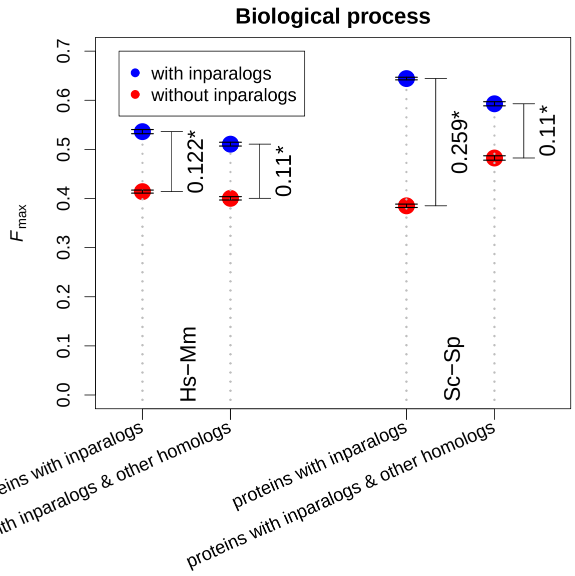


(b)

Figure S19: $F_{\max}$ values for proteins having inparalogs, proteins having inparalogs and other homologs (a), and proteins having both inparalogs and orthologs (b) in human-mouse and cerevisiae-pombe using biological process ontology (* indicates values are statistically significant).

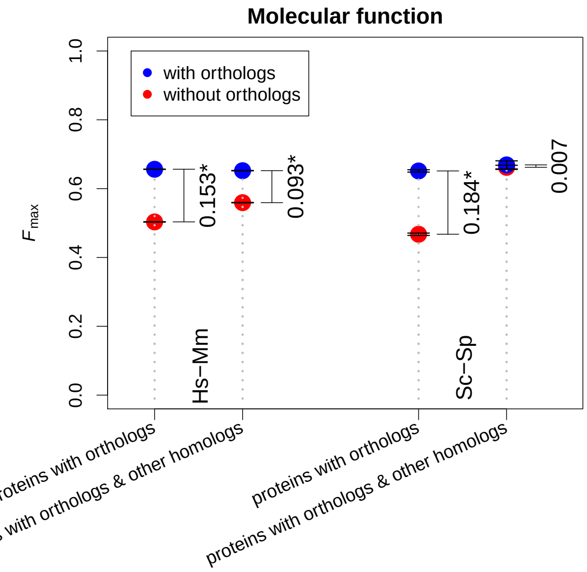


(a)

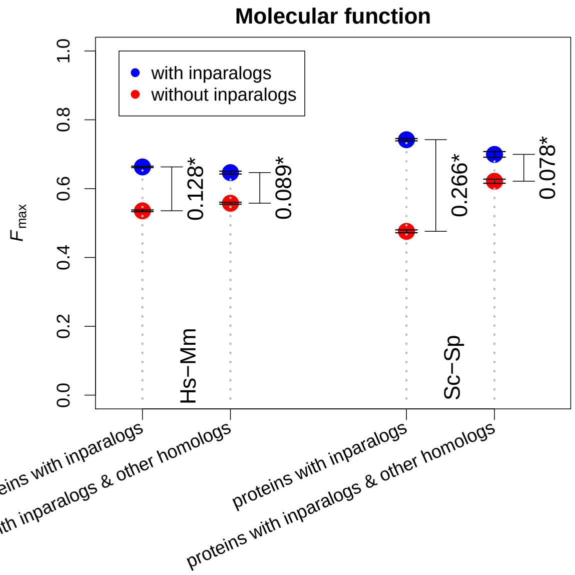


(b)

Figure S20: $F_{\max}$ values for proteins having inparalogs, proteins having inparalogs and other homologs (a), and proteins having both inparalogs and orthologs (b) in human-mouse and cerevisiae-pombe using molecular function ontology (* indicates values are statistically significant).

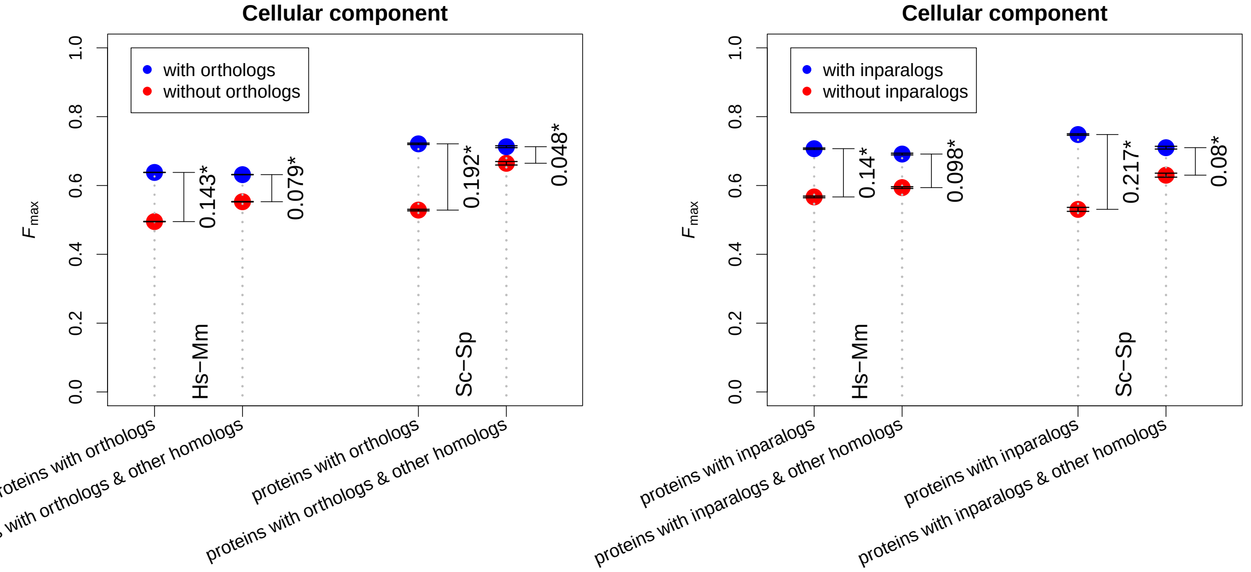


(a) (b)

Figure S21: $F_{\max}$ values for proteins having inparalogs, proteins having inparalogs and other homologs (a), and proteins having both inparalogs and orthologs (b) in human-mouse and cerevisiae-pombe using cellular context ontology (* indicates values are statistically significant).

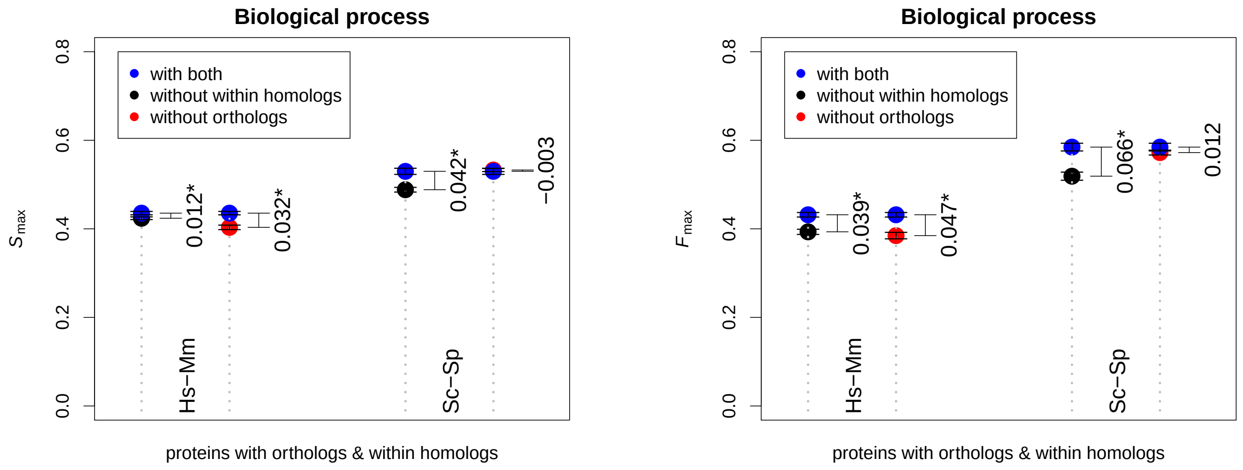


(a) (b)

Figure S22: $S_{\max}$ values for proteins both orthologs and within species homologs (a), Fmax values for proteins having both orthologs and within species homologs in human-mouse and cerevisiae-pombe using biological process ontology (* indicates values are statistically significant).

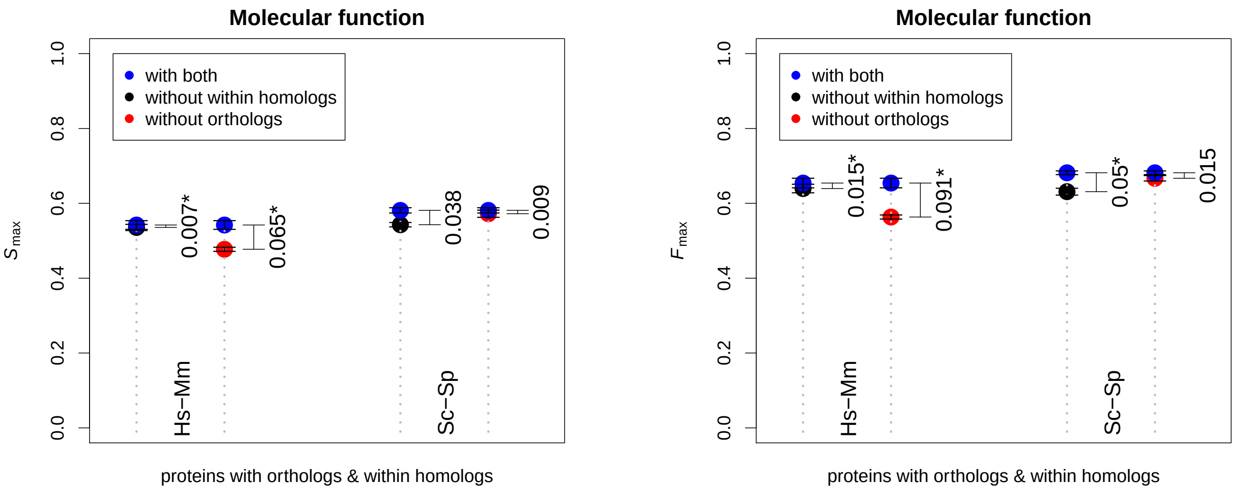


(a) (b)

Figure S23: $S_{\max}$ values for proteins both orthologs and within species homologs (a), Fmax values for proteins having both orthologs and within species homologs in human-mouse and cerevisiae-pombe using molecular function ontology (* indicates values are statistically significant).

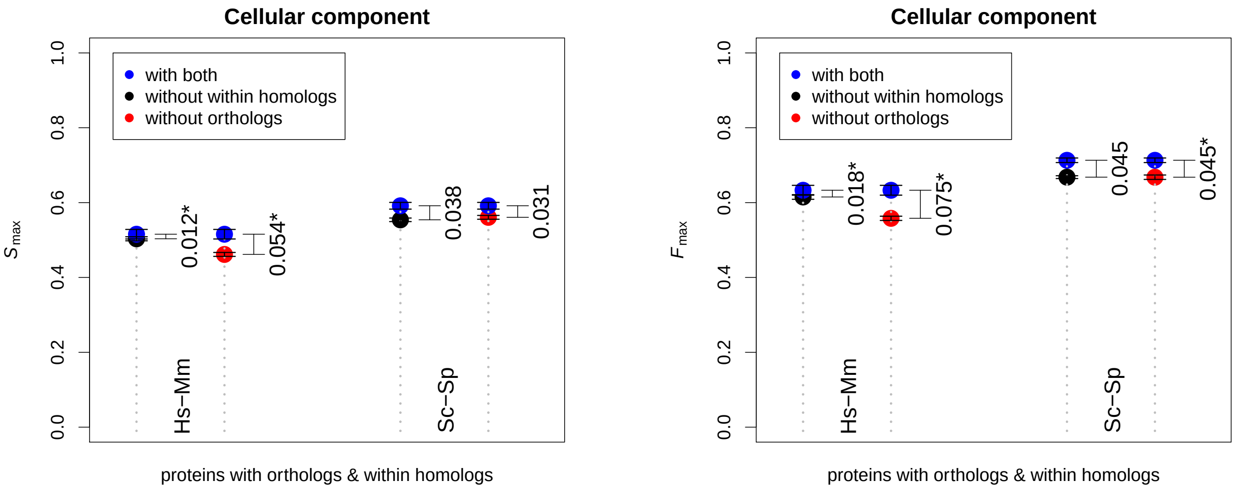


(a) (b)

Figure S24: $S_{\max}$ values for proteins both orthologs and within species homologs (a), Fmax values for proteins having both orthologs and within species homologs in human-mouse and cerevisiae-pombe using cellular context ontology (* indicates values are statistically significant).

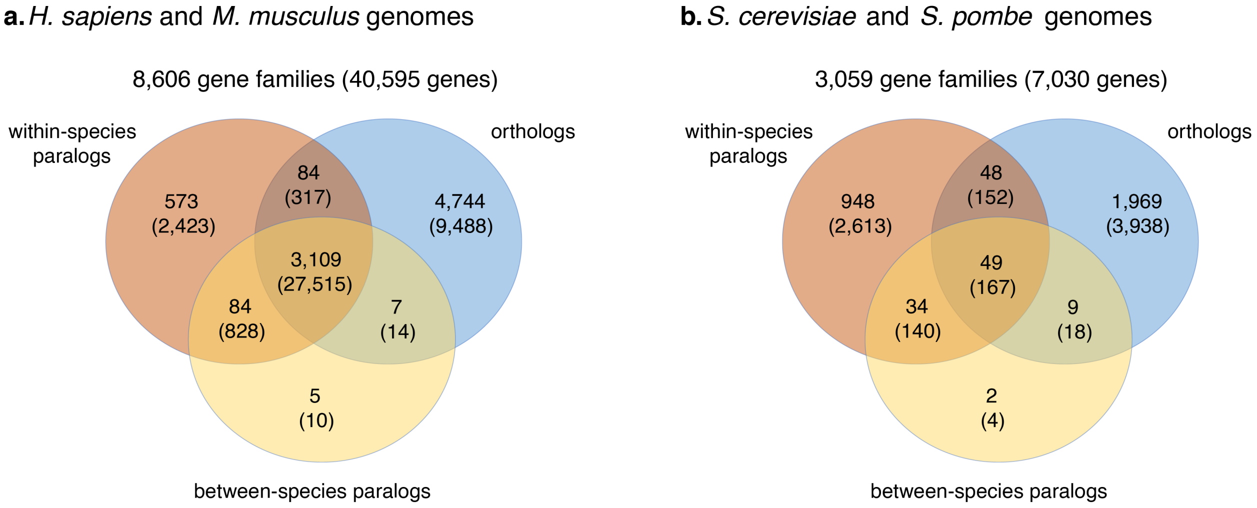


Figure S25: The numbers of gene families with different types of homologous relationships in them, where orthologs represent only one-to-one orthologs (Figure 3 in the main text considers all types of orthologs). The numbers in parentheses represent the counts of genes in the respective gene families. (a) gene families containing *H. sapiens* and *M. musculus* proteins, and (b) gene families containing *S. cerevisiae* and *S. pombe* proteins.

Table S1: Number of genes from human-mouse forming gene families, homologous pairs and number of families with different types of homologies.

| | | |
|---|---|---|
| Gene families (from gene trees) | 8686 | |
| genes making gene families | 41501 | (19,731 human + 21,770 mouse) |
| genes making homologous pairs | 40912 | (19,514 human + 21,398 mouse) |
| intersection (gene fams & hom pairs) | 40,884 | |
| protein coding genes | 44,603 | (22,376 human + 22,227 mouse) |
| intersection (p_coding genes & hom_genes) | 40,722 | |
| intersection (p_coding genes & gene families) | 40,741 | |
| total # of homologous pairs found in gene trees | 21,8254 | |
| ortholog pairs | 24,106 | |
| within species inparalog pairs | 33,741 | |
| within species outparalog pairs | 88,625 | |
| between specie outparalog pairs | 71,782 | |
| clusters containing homolgous pairs (at least 1 pair) | | |
| ortholog pairs | 8,606 | 99.07% |
| within inparalog pairs | 1,299 | 14.95% |
| between outparalog pairs | 3,205 | 36.89% |
| within outparalog pairs | 3,249 | 37.40% |
| both within inparalog and ortholog pairs | 1299 | 14.95% |
| both between homolog and within homolog | 3,850 | 44.32% |
| ortholog only families | 4,744 | 55.12% |

Table S2: Number of genes from cerevisiae-pombe forming gene families, homologous pairs and number of families with different types of homologies.

| | | |
|---|---|---|
| Gene families (from gene trees) | 3,061 | |
| genes making gene families | 7,335 | (4,042 yeast + 3,293 pombe) |
| genes making homologous pairs | 7,691 | (4,208 yeast + 3,483 pombe) |
| intersection (gene fams & hom pairs) | 7,334 | |
| protein coding genes | 11,839 | (6,693 yeast + 5,146 pombe) |
| intersection (p_coding genes & hom_genes) | 7,690 | |
| intersection (p_coding genes & gene families) | 3,061 | |
| total # of homologous pairs found in gene trees | 8,621 | |
| ortholog pairs | 3,249 | |
| within species inparalog pairs | 5,019 | |
| within species outparalog pairs | 202 | |
| between specie outparalog pairs | 151 | |
| clusters containing homolgous pairs (at least 1 pair) | | |
| ortholog pairs | 2,488 | 81.33% |
| within inparalog pairs | 999 | 32.60% |
| between outparalog pairs | 94 | 3.00% |
| within outparalog pairs | 130 | 4.20% |
| both within inparalog and ortholog pairs | 463 | 15.12% |
| both between homolog and within homolog | 1,079 | 35.27% |
| ortholog only families | 1,969 | 64.03% |

[6] Y. Jiang, T. R. Oron, W. T. Clark, A. R. Bankapur, D. D'Andrea, R. Lepore, C. S. Funk, I. Kahanda, K. M. Verspoor, A. Ben-Hur, C. E. Koo da, D. Penfold-Brown, D. Shasha, N. Youngs, R. Bonneau, A. Lin, S. M. Sahraeian, P. L. Martelli, G. Profiti, R. Casadio, R. Cao, Z. Zhong, J. Cheng, A. Altenhoff, N. Skunca, C. Dessimoz, T. Dogan, K. Hakala, S. Kaewphan, F. Mehryary, T. Salakoski, F. Ginter, H. Fang, B. Smithers, M. Oates, J. Gough, P. Toronen, P. Koskinen, L. Holm, C. T. Chen, W. L. Hsu, K. Bryson, D. Cozzetto, F. Minneci, D. T. Jones, S. Chapman, D. Bkc, I. K. Khan, D. Kihara, D. Ofer, N. Rappoport, A. Stern, E. Cibrian-Uhalte, P. Denny, R. E. Foulger, R. Hieta, D. Legge, R. C. Lovering, M. Magrane, A. N. Melidoni, P. Mutowo-Meullenet, K. Pichler, A. Shypitsyna, B. Li, P. Zakeri, S. ElShal, L. C. Tranchevent, S. Das, N. L. Dawson, D. Lee, J. G. Lees, I. Sillitoe, P. Bhat, T. Nepusz, A. E. Romero, R. Sasidharan, H. Yang, A. Paccanaro, J. Gillis, A. E. Sedeno-Cortes, P. Pavlidis, S. Feng, J. M. Cejuela,

T. Goldberg, T. Hamp, L. Richter, A. Salamov, T. Gabaldon, M. Marcet-Houben, F. Supek, Q. Gong, W. Ning, Y. Zhou, W. Tian, M. Falda, P. Fontana, E. Lavezzo, S. Toppo, C. Ferrari, M. Giollo, D. Piovesan, S. C. Tosatto, A. Del Pozo, J. M. Fernandez, P. Maietta, A. Valencia, M. L. Tress, A. Benso, S. Di Carlo, G. Politano, A. Savino, H. U. Rehman, M. Re, M. Mesiti, G. Valentini, J. W. Bargsten, A. D. van Dijk, B. Gemovic, S. Glisic, V. Perovic, V. Veljkovic, N. Veljkovic, E. S. D. C. Almeida, R. Z. Vencio, M. Sharan, J. Vogel, L. Kansakar, S. Zhang, S. Vucetic, Z. Wang, M. J. Sternberg, M. N. Wass, R. P. Huntley, M. J. Martin, C. O'Donovan, P. N. Robinson, Y. Moreau, A. Tramontano, P. C. Babbitt, S. E. Brenner, M. Linial, C. A. Orengo, B. Rost, C. S. Greene, S. D. Mooney, I. Friedberg, and P. Radivojac. An expanded evaluation of protein function prediction methods shows an improvement in accuracy. *Genome Biol*, 17(1):184, 2016.
